# A broadly conserved phytoplasma α/β-hydrolase effector, SGP-7, is a catalytically active esterase that hydrolyses strigolactone analogues and alters shoot architecture

**DOI:** 10.64898/2026.10.05.756913

**Authors:** Sneha Verma, Amit Yadav

## Abstract

Phytoplasmas are obligate, phloem-limited bacterial pathogens that reprogram host development through secreted effector proteins. The effectors characterized to date, including SAP11, SAP54/PHYL1, SAP05 and SWP1, act principally by binding and destabilizing host transcription factors. Although α/β-hydrolase-fold phytoplasma proteins with catalytic activity toward lipid substrates have recently been described, no phytoplasma effector has been shown to possess enzymatic activity toward a plant hormone. The effector RY378 from rice yellow dwarf (RYD) phytoplasma induces excessive tillering and dwarfing in rice and alters strigolactone– and auxin-responsive gene expression; its predicted α/β-hydrolase domain was proposed to mediate these effects, but the protein was never expressed, purified or biochemically characterized, leaving the mechanism of this effector class unresolved. From the closed genome of ‘*Ca.* P. sacchari’ isolate SCGS, associated with sugarcane grassy shoot disease, we identified SGP-7, an orthologue of RY378 that is conserved across at least seven 16Sr phytoplasma groups. SGP-7 adopts a canonical α/β-hydrolase fold with a Ser116– Asp229–His261 catalytic triad, and its predicted substrate pocket is comparable in volume to that of the *Arabidopsis* strigolactone receptor AtD14. Recombinant SGP-7 is a bona fide esterase with a clear preference for short-chain acyl esters, and hydrolyses both the fluorogenic strigolactone probe Yoshimulactone Green (apparent *K*_m_ 3.55 µM; *k*_cat_ 9.38 × 10⁻³ s⁻¹) and the strigolactone analogue *rac*-GR24. Both activities are abolished by substitution of the catalytic serine (SGP-7^S116A^). Transient expression of SGP-7 in *Nicotiana benthamiana* reduced plant height by 46%, increased axillary bud outgrowth approximately fourfold, and lowered transcript levels of the strigolactone-responsive marker *NbBRC1*, reproducing in a dicot host the architectural phenotype reported for RY378 in rice. Together, these results define the biochemical function of a conserved phytoplasma effector family and identify strigolactone-like molecules as direct enzymatic targets of a bacterial plant pathogen.

## 1. Introduction

Phytoplasmas (’*Candidatus* Phytoplasma’ spp.) are cell wall-less, phloem-restricted bacteria that have not yet been brought into axenic culture and that alternate between plant hosts and insect vectors. They are associated with disease in several hundred plant species worldwide and characteristically induce striking developmental abnormalities –witches’ broom, phyllody, virescence, dwarfism and excessive branching or tillering –that are far more conspicuous than the tissue damage they cause (Kirdat et al., 2023; Pedroncelli et al., 2026). Among them, ‘*Ca.* P. sacchari’ is associated with sugarcane grassy shoot disease, characterized by profuse tillering, stunted growth and proliferation of chlorotic shoots, which causes substantial yield losses in sugarcane-growing regions (Kirdat et al., 2021).

These symptoms are not a general consequence of infection but are directed by a small number of secreted effector proteins that enter host cells and reprogram development. The effectors characterized to date share a common mode of action: they bind host regulatory proteins and, in most cases, promote their degradation. SAP11 binds and destabilizes class II TCP transcription factors, producing crinkled leaves and stem proliferation (Pecher et al., 2019); SAP54/PHYL1 targets MADS-box transcription factors for ubiquitin-independent proteasomal degradation, converting flowers into leaf-like structures (MacLean et al., 2011); SAP05 hijacks the proteasomal Rpn10 subunit to degrade SPL and GATA transcription factors (Huang et al., 2021); and SWP1 destabilizes BRANCHED1 (BRC1) to promote witches’ broom (N. Wang et al., 2018). In each case, effector function is mediated through protein–protein interactions. No phytoplasma effector has yet been shown to act catalytically on a host small molecule.

Shoot architecture is governed by an integrated hormonal network in which strigolactones play a central role in suppressing axillary bud outgrowth and maintaining apical dominance (Al-Babili & Bouwmeester, 2015). Strigolactones are perceived by DWARF14 (D14), an α/β-hydrolase-fold receptor that is unusual in being both a receptor and an enzyme: it binds strigolactone, hydrolyses the D-ring through a Ser–Asp–His catalytic triad, and undergoes a hydrolysis-dependent conformational change that permits recruitment of the F-box protein MAX2/D3 and consequent degradation of the SMXL6/7/8 repressors (Hamiaux et al., 2012; Yao et al., 2016). Because SMXL6/7/8 repress *BRC1* transcription, strigolactone signaling normally elevates *BRC1* and thereby reinforces bud dormancy (L. Wang et al., 2015). Mutants defective in strigolactone biosynthesis or signaling are dwarfed and highly branched (Gomez-Roldan et al., 2008; Umehara et al., 2008), a phenotype that closely resembles phytoplasma-induced witches’ broom and has long invited the hypothesis that phytoplasmas interfere with this pathway.

Direct evidence for this hypothesis has recently emerged. Zhang and colleagues sequenced the genome of rice yellow dwarf (RYD) phytoplasma strain HN2022 and identified RY378 as a virulence effector: transient expression in *N. benthamiana* produced witches’ broom-like symptoms, while stable overexpression in rice increased tillering by approximately 350% and reduced plant height by one third (Zhang et al., 2024). Transcriptome and RT-qPCR analyses of RY378-overexpressing rice showed altered expression of strigolactone– and auxin-pathway genes, including *D3*, *IPA1*, *T20*, *IAA6*, *PIN1* and *PIN5b* (Zhang et al., 2024). Critically, the same study noted that RY378 carries a predicted α/β-hydrolase domain and proposed this domain as the structural basis of its function, but the protein was not produced recombinantly, no substrate was tested, no catalytic residue was mutated and no enzymatic parameter was measured. The molecular function of this effector class therefore remained an explicit and untested hypothesis. Zhang and colleagues further reported that they could identify RY378 homologues only in SCWL1, from the same 16SrXI group, and concluded that RY378 is lineage-restricted and may represent a unique pathogenic mechanism (Zhang et al., 2024).

Characterisation of other phytoplasma genomes has identified additional α/β-hydrolase-domain proteins with enzymatic activity. A lipase-like enzyme from potato purple top phytoplasma (PPT-lipase) is a predicted monoacylglycerol lipase with a conserved catalytic triad that reduces neutral lipid levels in heterologous expression systems and localizes intracellularly; it is proposed to process host lipid intermediates during infection (Inaba et al., 2026). That work established that phytoplasmas encode catalytically competent α/β-hydrolases, and that such enzymes may diversify toward distinct metabolic substrates. Whether α/β-hydrolase effectors from other phytoplasma lineages might instead act on signaling molecules, in particular the plant hormones that govern meristematic development, has remained unexplored.

This gap can be addressed directly. If the α/β-hydrolase domain of this effector family is catalytically competent, and if its substrate preference resembles that of the strigolactone receptors it structurally mimics, the effector would represent a mechanistically distinct class of phytoplasma virulence factor that acts on a hormone molecule rather than on a host protein. In this study, SGP-7, the ‘*Ca.* P. sacchari’ SCGS orthologue of RY378, is identified from a closed and polished genome assembly; its distribution across phytoplasma lineages is defined; its fold, catalytic triad and predicted substrate pocket are determined; the recombinant protein and an active-site mutant are expressed and purified; and its ability to hydrolyze strigolactone analogues is tested directly. Transient expression of SGP-7 in *N. benthamiana* is then used to determine whether it reproduces the architectural changes reported for RY378 in rice.

## 2. Results

### 2.1. A closed genome assembly for ‘*Ca.* P. sacchari’ SCGS defines a repertoire of seven candidate effectors

To establish a complete gene inventory for effector discovery, the previously reported draft assembly of ‘*Ca.* P. sacchari’ isolate SCGS (29 scaffolds, 505.2 kb; GCA_009268105.1; Kirdat et al., 2021) was re-sequenced using Oxford Nanopore long reads and polished with existing Illumina data. The resulting assembly comprises a single circular chromosome of 562,492 bp (CP176574.1) and a 2,949-bp plasmid (CP176575.1), giving a total assembly size of 565,441 bp, with an overall G+C content of 20.5%. Annotation identified 461 protein-coding sequences, including 110 hypothetical proteins, together with 27 tRNA genes, two complete rRNA operons, two non-coding RNAs and one pseudogene (Table 1; Fig. 1A). Genome completeness was estimated at 98% by BUSCO (phytoplasma database) and 94.18% by CheckM, with 0.58% contamination. KEGG annotation assigned 417 genes to functional categories, and COG classification assigned 327 proteins, with the largest categories corresponding to genetic information processing and translation, respectively (Fig. 1B, C).

**Figure 1.**
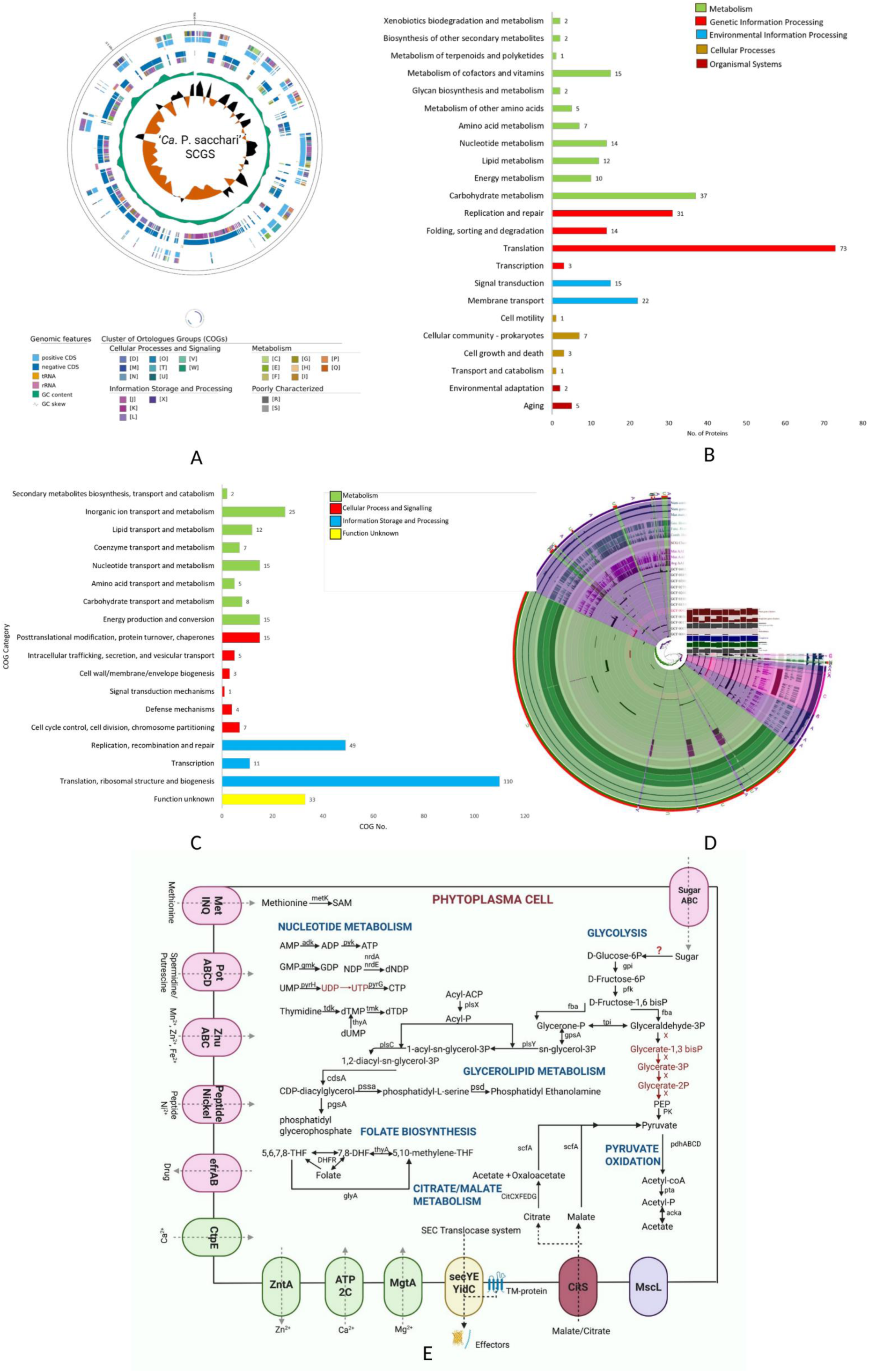
Genomic features of the closed ‘*Candidatus* Phytoplasma sacchari’ isolate SCGS assembly. **(A)** Circular map of the SCGS chromosome showing, from the outside inwards, genomic features, Clusters of Orthologous Groups (COG) assignments and metabolic categories. **(B)** Distribution of genes across KEGG functional categories, colored by top-level pathway class. **(C)** Distribution of proteins across COG functional categories. **(D)** Pan-genome analysis of SCGS together with 11 complete phytoplasma reference genomes, showing core, accessory and strain-specific gene clusters. **(E)** Reconstruction of central metabolic and transport capacity in the SCGS cell, including the Sec-dependent secretion system, nucleotide and glycerolipid metabolism, folate biosynthesis and predicted transporters.

**Table 1.** Comparison of genome features between ‘*Candidatus* Phytoplasma sacchari’ isolate SCGS and rice yellow dwarf (RYD) phytoplasma strain HN2022.

| Feature | 'Ca. P. sacchari' SCGS | RYD phytoplasma HN2022 |
| --- | --- | --- |
| 16Sr group | 16SrXI-B | 16SrXI-B |
| Chromosome size (bp) | 562,492 | 511,268 |
| Chromosome organisation | Circular | Circular |
| Extrachromosomal element | 1 plasmid (2,949 bp) | None |
| Total assembly size (bp) | 565,441 | 511,268 |
| G+C content (%) | 20.5 | 20.25 |
| Coding density (%) | 76.62 | 79.3 |
| Protein-coding genes (CDS) | 461 | 423 |
| Hypothetical proteins | 110 | 92 |
| tRNA genes | 27 | 24 |
| rRNA operons | 2 | 2 |
| Total genes | 496 | 455 |
| Sequencing platform | Oxford Nanopore + Illumina | PacBio HiFi |
| Genome accession | CP176574.1 (chromosome);<br>CP176575.1 (plasmid) | CP116038.1 |
| Reference | This study | Zhang et al., 2024 |

Pan-genome analysis against 11 complete reference phytoplasma genomes identified 2,304 gene clusters, comprising 146 core, 688 accessory and 1,470 unique clusters (Fig. 1D). Consistent with the reduced genomes characteristic of phytoplasmas (Oshima et al., 2004), SCGS lacks *de novo* nucleotide biosynthesis and contains only a partial glycolytic pathway. In contrast, it retains an intact Sec-dependent secretion system, including *secY*, *secE*, *secA*, signal peptidase, *dnaJ*, *dnaK*, *groES*, *groEL*, *ffh* and *ftsY*, as well as 30 predicted transporters, including a complete ABC-type multiple-sugar transport system (Fig. 1E).

The combination of a closed genome, a substantial fraction of hypothetical proteins and an intact secretion machinery provided a defined framework for systematic effector discovery. Among the 461 predicted proteins, seven candidates were prioritized on the basis of features associated with phytoplasma effectors, including secretion-related signals, subcellular localization predictions and sequence characteristics. These candidates were designated SGP-1 to SGP-7.

### 2.2. SGP-7 is a broadly conserved, secreted phytoplasma α/β-hydrolase effector

The seven candidate secreted effectors were identified in the SCGS proteome on the basis of a predicted N-terminal signal peptide (SignalP 6.0) and the absence of transmembrane helices in the mature protein (TMHMM 2.0). Except for SGP-7, which was annotated as an α/β-hydrolase, all candidates were annotated as hypothetical proteins; only SGP-3 carried a predicted nuclear localization signal (Table 2). To assess their distribution, a curated secretome database was assembled from 288 publicly available phytoplasma genomes and searched for homologues using BLASTP.

**Table 2.** Candidate secreted effectors encoded by the ‘*Candidatus* Phytoplasma sacchari’ isolate SCGS genome and their distribution across phytoplasma lineages.

| Effector | Accession | Length (aa) | Homologues (n) | 16Sr group distribution | Distribution class | Predicted NLS |
| --- | --- | --- | --- | --- | --- | --- |
| SGP-1 | XXZ63711.1 | 214 | 1 | 16SrXI, 16SrXIV | Narrowly distributed | No |
| SGP-2 | XXZ63718.1 | 104 | 0 | 16SrXI | 'Ca. P. sacchari'-specific | No |
| SGP-3 | XXZ63746.1 | 144 | 0 | 16SrXI | 'Ca. P. sacchari'-specific | Yes |
| SGP-4 | XXZ63857.1 | 378 | 10 | 16SrII, III, VII, XI, XIV, XXI | Broadly conserved | No |
| SGP-5 | XXZ63998.1 | 193 | 0 | 16SrXI | 'Ca. P. sacchari'-specific | No |
| SGP-6 | XXZ64085.1 | 261 | 5 | 16SrI, XI, XIV | Narrowly distributed | No |
| SGP-7 | XXZ64048.1 | 320 | 15 | 16SrI, II, III, VII, X, XI, XIV | Broadly conserved ($\alpha/\beta$-hydrolase) | No |

The seven candidates showed distinct distribution patterns. SGP-2, SGP-3 and SGP-5 had no homologues outside ‘*Ca.* P. sacchari’, indicating lineage-specific distribution. SGP-1 and SGP-6 were narrowly distributed, with one and five homologues respectively, confined to two or three phytoplasma groups. SGP-4 was more conserved, with 10 homologues across the 16SrII, III, VII, XI, XIV and XXI groups. SGP-7 showed the broadest distribution, with 15 homologues across seven groups: 16SrI, II, III, VII, X, XI and XIV (Table 2). These groups include phytoplasmas infecting both monocot and dicot hosts.

SGP-7 was selected for functional analysis because it was the only candidate carrying both a predicted N-terminal signal peptide and a predicted catalytic domain (α/β-hydrolase fold), a combination that distinguishes it from other characterized phytoplasma α/β-hydrolases. Although phytoplasma α/β-hydrolases have been identified, the characterized examples are intracellular enzymes acting on metabolic substrates (Inaba et al., 2026). No phytoplasma α/β– hydrolase has previously been shown to function as a secreted effector targeting plant hormones or other host developmental signaling molecules.

The broad distribution of SGP-7 across seven phytoplasma groups also contrasts with the proposed lineage restriction of its RY378 orthologue. Zhang and colleagues detected RY378 homologues only in SCWL1, a second member of the 16SrXI group, and inferred that the effector was specific to 16SrXI and might mediate a lineage-restricted pathogenic mechanism (Zhang et al., 2024). In contrast, the larger, purpose-built secretome database used here identified 15 homologues across seven 16Sr groups. This difference may reflect database scope, since searches limited to annotated reference proteomes can miss effectors present in draft genomes or proteins annotated as hypothetical. SGP-7 is therefore best regarded as a representative of a conserved, secreted phytoplasma effector family rather than a strain-specific virulence factor, suggesting that its biochemical properties and host targets may be broadly relevant to phytoplasma pathogenicity.

### 2.3. SGP-7 adopts an α/β-hydrolase fold with a Ser116–Asp229–His261 catalytic triad and a D14-like substrate pocket

SGP-7 is a 320-residue protein with a 33-residue N-terminal signal peptide, giving a mature protein of 287 residues, and is annotated as an α/β-hydrolase (Fig. 2A). Throughout this study, residues are numbered according to the mature protein. Multiple sequence alignment against α/β-hydrolase homologues retrieved from the ESTHER database identified a candidate catalytic triad comprising Ser116, Asp229 and His261, with the nucleophilic serine positioned within a canonical G-X-S-X-G motif (G114-H115-S116-L117-G118) (Supplementary Figure S3). Alignment with the plant strigolactone receptors AtD14, OsD14 and DAD2 showed that all three catalytic positions are structurally equivalent to the receptor triad (Supplementary Figure S7).

**Figure 2.**
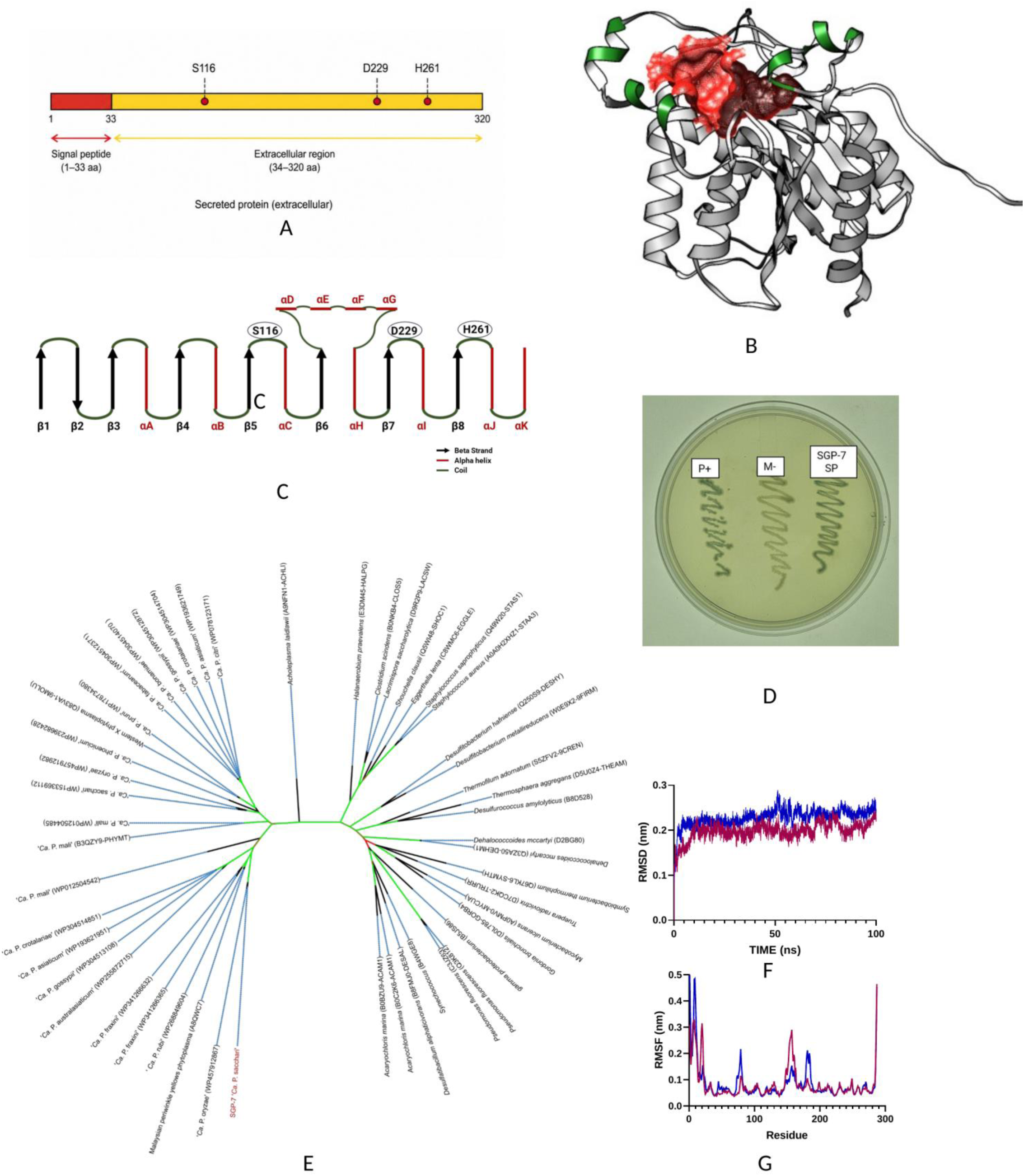
SGP-7 is a secreted α/β-hydrolase with a canonical catalytic triad and a stable predicted fold. **(A)** Domain architecture of SGP-7, showing the 33-residue N-terminal signal peptide, the mature extracellular region (residues 34–320 in full-length numbering) and the positions of the catalytic residues Ser116, Asp229 and His261 (mature-protein numbering). **(B)** Principal surface cavity of SGP-7 predicted by CASTp 3.0, shown in red on the AlphaFold model; the cavity has a solvent-accessible area of 231.7 Å² and a solvent-accessible volume of 167.3 Å³ and is lined by 24 residues including Ser116 and His261. **(C)** Topology diagram of the predicted SGP-7 fold, showing eight β-strands and eleven α-helices, with the catalytic residues located in the loops connecting β5–αC, β7–αI and β8–αJ. **(D)** Alkaline phosphatase (PhoA) secretion assay on BCIP-containing medium. P+, full-length *E. coli* PhoA (positive control); N−, signal-peptide-less mPhoA (negative control); SGP-7 SP, mPhoA fused to the predicted SGP-7 signal peptide. Blue colonies indicate periplasmic export. **(E)** Maximum-likelihood phylogeny of SGP-7 with the top 30 ESTHER-database α/β-hydrolases and the top 20 phytoplasma homologues from 19 RefSeq reference genomes, inferred with IQ-TREE under Q.PFAM+R5 with 1,000 bootstrap replicates. Branch color indicates bootstrap support (green, high; red, low). **(F)** Backbone RMSD of apo SGP-7 (blue) and the SGP-7– GR24 complex (magenta) over the 100 ns simulation. **(G)** Per-residue RMSF for the same two systems.

The AlphaFold-predicted model of mature SGP-7 showed good stereochemical quality, with 97.54% of residues in the favored regions and 2.46% in the additionally allowed regions of the Ramachandran plot, and no residues in disallowed regions (Supplementary Figure S2), supporting the reliability of the predicted structure. The model comprises eight β-strands and eleven α-helices connected by loops, with the second β-strand arranged antiparallel to the remainder. The structure is organised into a catalytic domain and a lid domain formed by four α-helices located between β6 and αH. The catalytic residues Ser116, Asp229 and His261 lie in the loops connecting β5–αC, β7–αI and β8–αJ respectively, corresponding to the canonical positions for this fold (Fig. 2C). The functional competence of the predicted signal peptide was confirmed experimentally using an alkaline phosphatase (PhoA) fusion assay, in which the SGP-7 signal peptide directed export of a signal-peptide-less PhoA to the periplasm and produced blue colonies on BCIP-containing medium (Fig. 2D).

CASTp analysis identified a principal surface pocket with a solvent-accessible surface area of 231.7 Å² and a solvent-accessible volume of 167.3 Å³, lined by 24 residues including both Ser116 and His261 (Fig. 2B). Applying the same analysis under identical settings to the *Arabidopsis* strigolactone receptor AtD14 (PDB 4IH4) returned a pocket of 321.6 Å² and 166.9 Å³ (Supplementary Table S1). The predicted SGP-7 cavity is therefore closely comparable in volume to the strigolactone-binding pocket of AtD14, while being somewhat more enclosed, indicating that SGP-7 possesses a cavity of appropriate dimensions to accommodate a strigolactone-sized ligand.

To assess whether the predicted model represents a stable fold, the structure was subjected to a 100 ns all-atom molecular dynamics simulation. Backbone RMSD stabilized after approximately 10 ns and thereafter fluctuated between 0.16 and 0.25 nm (mean 0.20 nm) (Fig. 2F). Per-residue RMSF averaged 0.08 nm across the 287 residues and remained below 0.1 nm for the large majority of the chain, with larger excursions (up to 0.47 nm) confined to the termini and to a small number of surface loops (Fig. 2G). The radius of gyration was stable at 1.84–1.89 nm (mean 1.86 nm) and the solvent-accessible surface area fluctuated around 133–154 nm² (mean 143 nm²) without sustained drift, indicating that overall compactness and solvent exposure were maintained (Supplementary Figure S5A, B). Collectively, these parameters indicate that the SGP-7 model remained compact and conformationally stable throughout the simulation. Maximum-likelihood phylogenetic analysis of SGP-7 together with ESTHER-derived α/β-hydrolases and homologues from 19 reference phytoplasma genomes placed SGP-7 within a well-supported phytoplasma-specific clade, clearly separated from the characterized α/β– hydrolase families represented in ESTHER (Fig. 2E).

### 2.4. Recombinant SGP-7 and the active-site mutant SGP-7^S116A^ can be purified as stable monomers

To test the predicted catalytic function directly, the mature SGP-7 coding sequence was codon-optimized, cloned with a C-terminal hexa-histidine tag and expressed in *Escherichia coli* BL21(DE3). A catalytically inactive variant, SGP-7^S116A^, was generated by site-directed mutagenesis of the nucleophilic serine and verified by Sanger sequencing. Both proteins were recovered from the soluble fraction and purified by immobilized metal affinity chromatography, yielding a single dominant band at approximately 34 kDa (**Fig. 3A**). Size-exclusion chromatography on a Superdex 200 Increase column resolved a single symmetrical peak eluting at 16–17 mL, consistent with a monomeric ∼35 kDa species, and SDS-PAGE of the peak fractions confirmed the presence of purified SGP-7 (**Fig. 3B**; **Supplementary Figure S4**). The identity of both proteins was confirmed by anti-His immunoblotting (**Fig. 3C**). Intact mass spectrometry of wild-type SGP-7 gave an observed mass of 34,541.7 Da, in close agreement with the mass of 34,542.3 Da predicted for the tagged construct and indicating retention of the N-terminal methionine; peptide-level analysis of the trypsin-digested protein identified 22 unique peptides covering 76.6% of the sequence.

**Figure 3.**
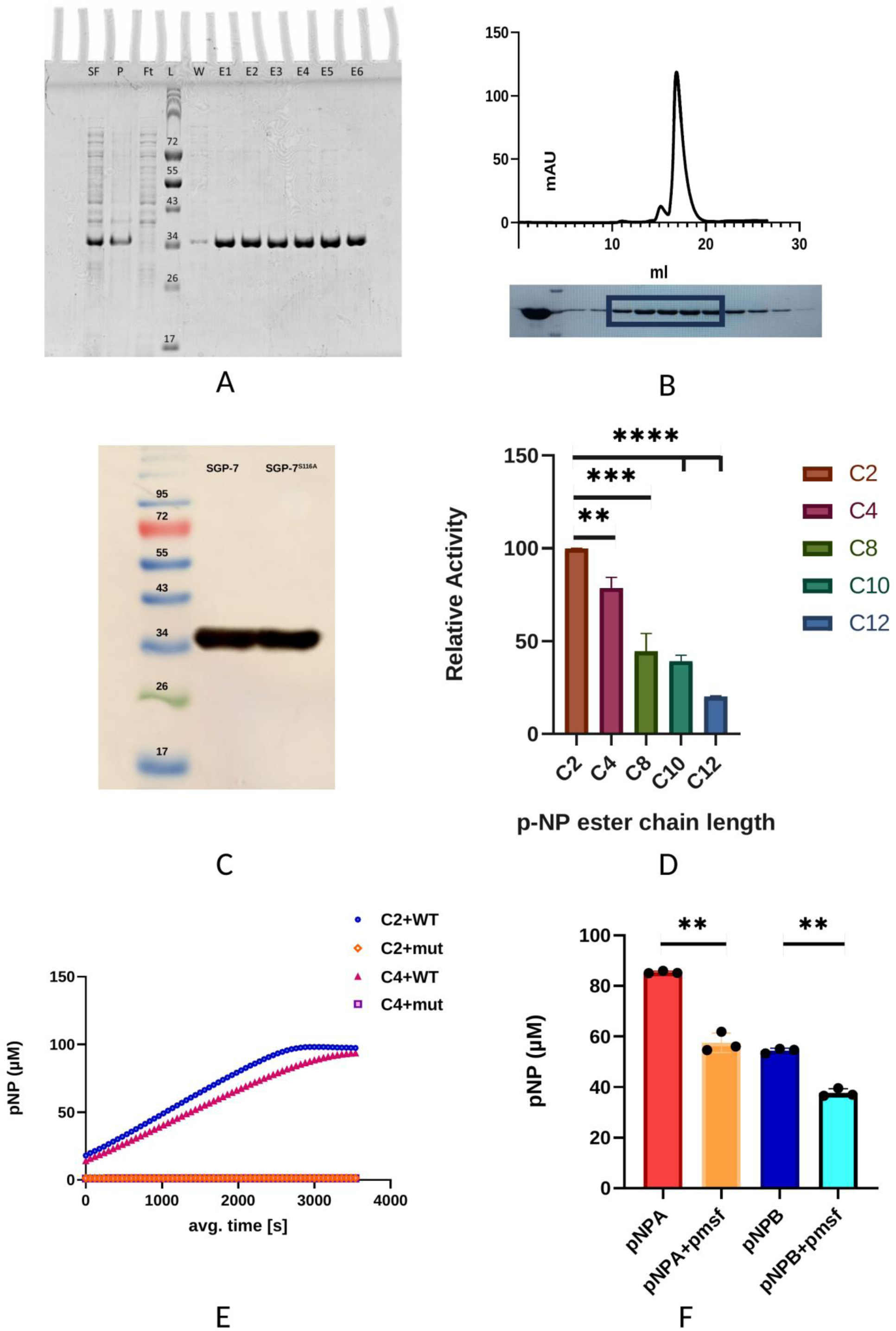
Recombinant SGP-7 is a monomeric, Ser116-dependent esterase with a preference for short-chain acyl esters. **(A)** SDS-PAGE of immobilised metal affinity chromatography fractions. SF, soluble fraction; P, pellet; Ft, flow-through; L, ladder; W, wash; E1–E6, elution fractions. **(B)** Size-exclusion chromatogram on a Superdex 200 Increase 10/300 GL column, with SDS-PAGE of the peak fractions shown beneath; the single symmetrical peak at 16–17 mL is consistent with a monomer. **(C)** Anti-6×His immunoblot of purified SGP-7 and SGP-7^S116A^. **(D)** Relative activity of SGP-7 against *p*-nitrophenyl esters of increasing acyl chain length (C2–C12). Activity is normalized within each assay to C2 = 100%; bars show mean ± SD of three independent assays, each with four technical replicates. **(E)** Time course of *p*-nitrophenol release from 100 µM C2 and C4 substrates by wild-type SGP-7 and SGP-7^S116A^ (mut). Each point is the mean of eight technical replicates. **(F)** Effect of 1 mM PMSF on *p*-nitrophenol release from 1 mM *p*-nitrophenyl acetate (pNPA) and *p*-nitrophenyl butyrate (pNPB) after 30 min at 30 °C; mean ± SD of three independent assays. Note that the substrate concentration in this panel is ten-fold higher than in panels D and E. \*\**P* < 0.01, P *< 0.001,* P *< 0.0001 (unpaired two-tailed Student’s* t*-test).

Importantly, SGP-7^S116A^ was expressed, purified and eluted indistinguishably from the wild-type protein, indicating that the substitution does not substantially affect protein folding or stability, and therefore that any loss of activity results from removal of the catalytic nucleophile rather than from misfolding.

### 2.5. SGP-7 is an active esterase with a marked preference for short-chain acyl esters

The catalytic activity of purified SGP-7 was first assessed using a homologous series of *p*-nitrophenyl esters of increasing acyl chain length. SGP-7 was most active toward *p*-nitrophenyl acetate (C2), which was set to 100% relative activity within each independent assay. Activity declined progressively with increasing chain length, to 78.6 ± 5.6% for *p*-nitrophenyl butyrate (C4), 44.5 ± 9.7% for octanoate (C8), 39.3 ± 3.0% for decanoate (C10) and 20.1 ± 0.4% for dodecanoate (C12) (mean ± SD, *n* = 3 independent assays) (**Fig. 3D**). This monotonic decline across the C2–C12 series is consistent with esterase activity and a preference for short-chain substrates rather than with lipase activity.

Time-resolved assays confirmed that hydrolysis is enzymatic and strictly dependent on Ser116. Wild-type SGP-7 progressively hydrolysed both C2 and C4 substrates over one hour, approaching complete conversion of the 100 µM substrates, whereas SGP-7^S116A^ produced no detectable *p*-nitrophenol above background throughout the time course (**Fig. 3E**). Treatment with the serine hydrolase inhibitor phenylmethylsulfonyl fluoride (PMSF) at 1 mM reduced *p*-nitrophenol release by 32.6% for C2 and 30.7% for C4 (*P* < 0.001 for both) (**Fig. 3F**), consistent with a mechanism involving the active-site serine. The partial nature of this inhibition is considered in the Discussion.

### 2.6. SGP-7 hydrolyses the strigolactone probe Yoshimulactone Green with receptor-like kinetics

The structural correspondence between SGP-7 and D14-family strigolactone receptors –a shared fold, an equivalent catalytic triad and a substrate pocket of comparable volume –prompted an assessment of whether SGP-7 acts on strigolactone-like molecules (**Supplementary Figure S8A**).

Yoshimulactone Green (YLG), a fluorogenic strigolactone analogue whose hydrolysis at the D-ring releases fluorescein and which is widely used to measure strigolactone receptor activity, was used first. Wild-type SGP-7 readily hydrolyzed YLG, producing a rapid increase in fluorescein fluorescence that approached a plateau at approximately 3,000 s, whereas SGP-7^S116A^ generated only background fluorescence throughout the assay (**Fig. 5C**). Hydrolysis of YLG by SGP-7 is therefore strictly dependent on the catalytic serine.

**Figure 4.**
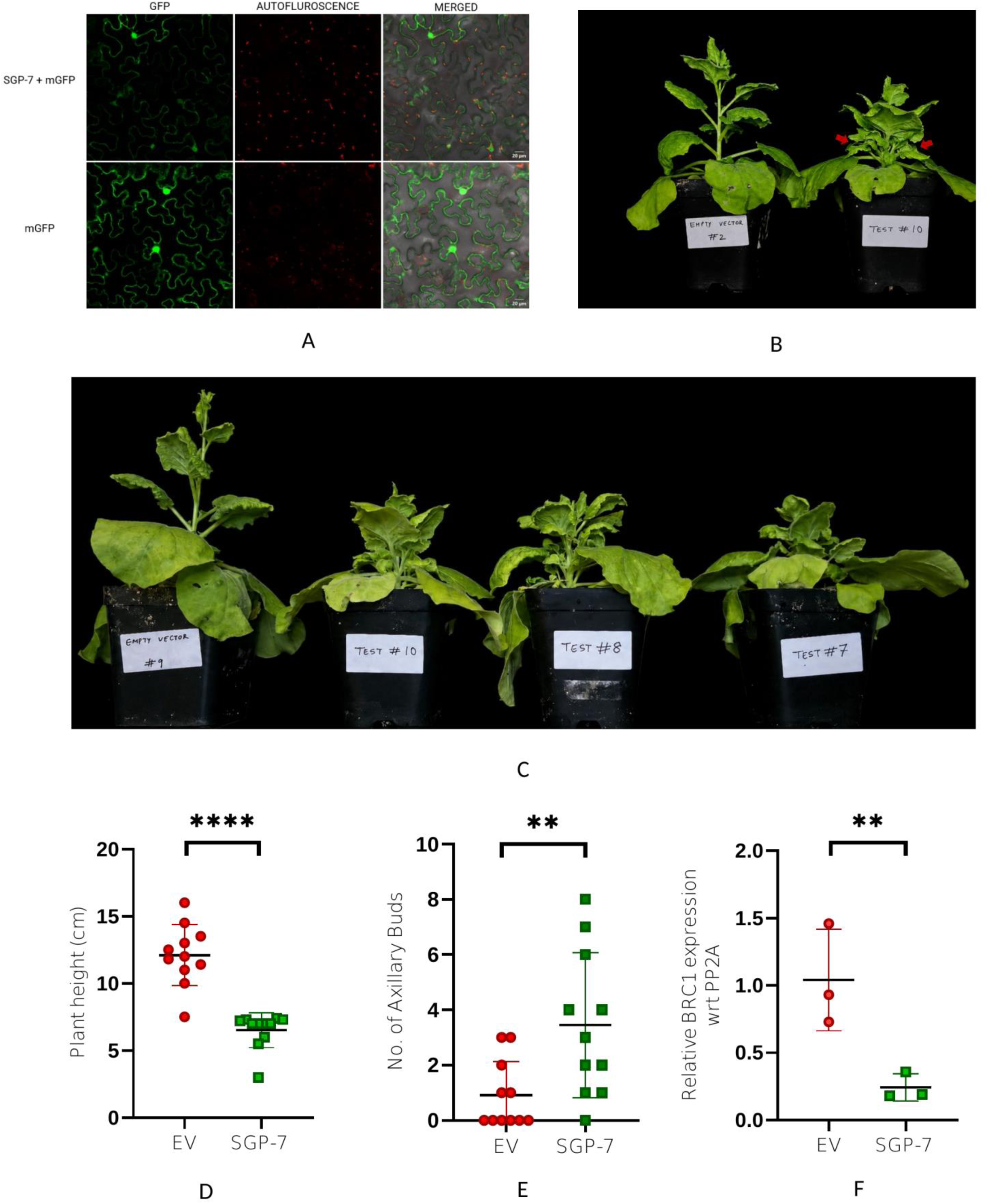
Transient expression of SGP-7 in *Nicotiana benthamiana* reduces stature and releases axillary buds. **(A)** Confocal imaging of *N. benthamiana* epidermal cells expressing SGP-7–mGFP or free mGFP, showing GFP fluorescence, chlorophyll autofluorescence and merged channels. Both constructs give a nucleocytoplasmic distribution. **(B)** Representative empty-vector and SGP-7-expressing plants at 21 days post-infiltration; red arrows indicate axillary buds. **(C)** Representative plants illustrating the reduction in stature in SGP-7-expressing plants relative to the empty-vector control. **(D)** Plant height at 21 days post-infiltration (*n* = 11 plants per treatment, each plant one biological replicate). **(E)** Number of axillary buds exceeding 2 cm per plant at 21 days post-infiltration (*n* = 11 per treatment); significance was assessed using a two-tailed Mann– Whitney U test. **(F)** Relative *NbBRC1* transcript abundance in axillary bud tissue, normalized to *PP2A* and expressed relative to the empty-vector control (*n* = 3 biological replicates, each with three technical replicates); Unless otherwise indicated, significance assessed by unpaired two-tailed Student’s *t*-test on Δ*C*_t_ values. Bars show mean ± SD. \*\**P* < 0.01, *P* < 0.0001. EV, empty vector.

**Figure 5.**
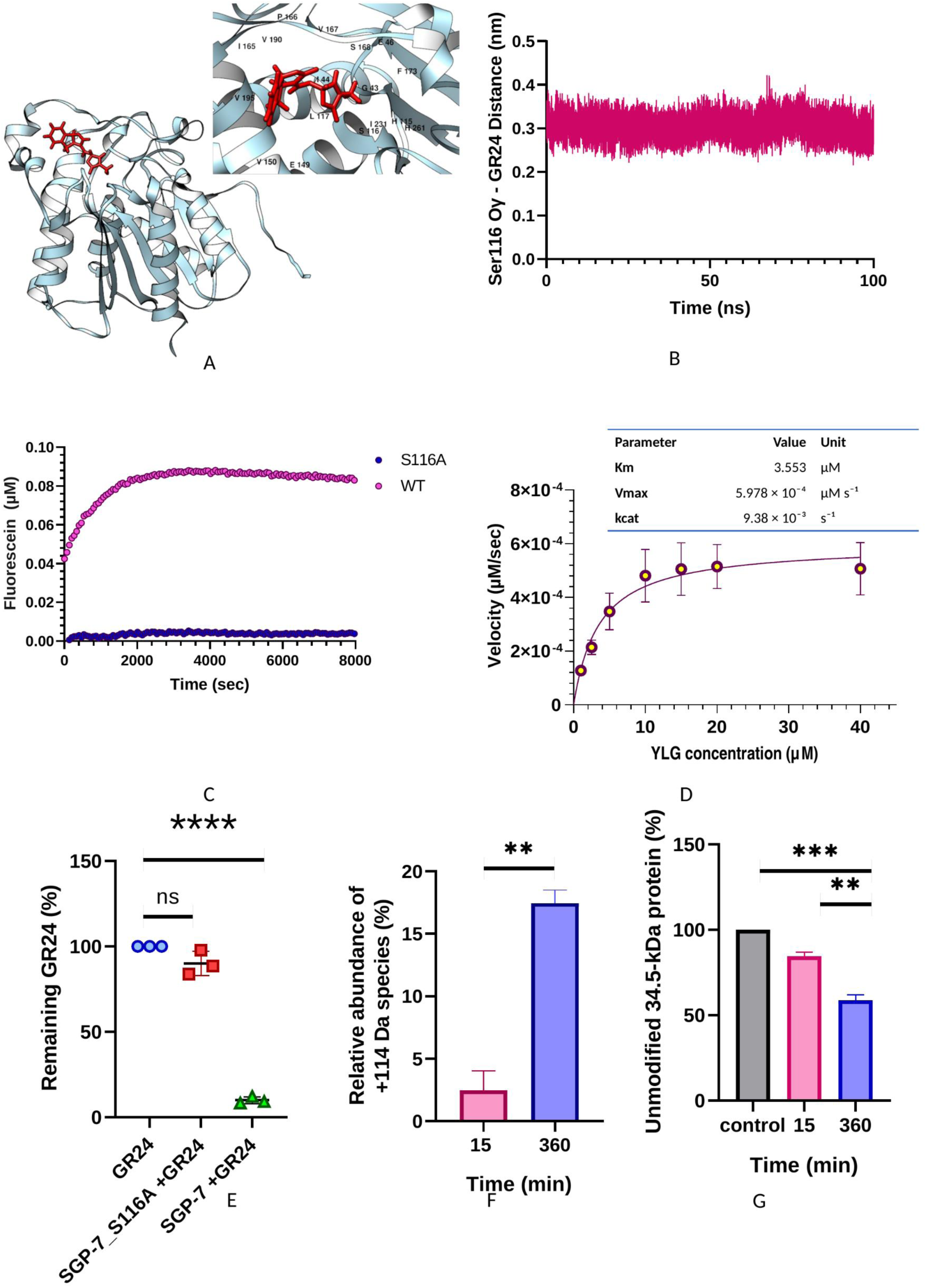
SGP-7 hydrolyzes the strigolactone analogs YLG and *rac*-GR24 in a Ser116-dependent manner. **(A)** GR24 docked into the predicted binding pocket of SGP-7, with the inset showing the residues surrounding the ligand. **(B)** Minimum distance between the Ser116 Oγ atom and GR24 across the 100 ns molecular dynamics simulation (50,001 frames; mean 0.31 nm). **(C)** Time course of fluorescein release from 1 µM Yoshimulactone Green by wild-type SGP-7 (magenta) and SGP-7^S116A^ (blue); eight technical replicates. **(D)** Michaelis–Menten analysis of YLG hydrolysis by SGP-7 across 1–40 µM substrate, with fitted parameters inset (three biological replicates, four technical replicates each; mean ± SD). **(E)** GR24 remaining after 30 min incubation, quantified by HPLC and expressed as a percentage of the substrate-only control (three biological replicates, three technical replicates each). **(F)** Relative abundance of the +114 Da modified species of SGP-7 after incubation with GR24 for 15 and 360 min (three technical replicates per time point). **(G)** Abundance of the unmodified 34.5-kDa species at the same time points, relative to the untreated control. Bars show mean ± SD. ns, not significant; \*\**P* < 0.01, P *< 0.001,* P* < 0.0001.

Steady-state kinetic analysis was performed across YLG concentrations from 1 to 40 µM. Initial velocities determined from the linear phase of each progress curve increased with substrate concentration and approached saturation, and the data fitted the Michaelis–Menten equation, yielding an apparent *K*_m_ of 3.55 µM, a *V*_max_ of 5.98 × 10⁻⁴ µM s⁻¹ and a *k*_cat_ of 9.38 × 10⁻³ s⁻¹ (**Fig. 5D**). These parameters fall within the range reported for plant strigolactone receptors: the apparent *K*_m_ is approximately twofold higher than the value of 1.84 µM reported for AtD14 with the same probe (Nakamura et al., 2019), while the turnover number is of the same low order reported for D14-family proteins, which behave kinetically as slow, near-single-turnover enzymes rather than as efficient metabolic hydrolases. SGP-7 therefore represents a lower-affinity, slow-turnover hydrolase of a strigolactone analogue, with kinetic properties resembling those of a receptor rather than those of a catabolic enzyme.

### 2.7. SGP-7 hydrolyses *rac*-GR24 and undergoes a GR24-dependent covalent modification

To determine whether SGP-7 also acts on the synthetic strigolactone analogue *rac*-GR24, purified wild-type SGP-7 and the S116A mutant were incubated with *rac*-GR24 and the remaining substrate was quantified by HPLC against a substrate-only control. Incubation with wild-type SGP-7 reduced the remaining GR24 to 9.9 ± 2.0% of the control, corresponding to approximately 90% hydrolysis of the substrate (*P* < 0.0001). In contrast, 90.1 ± 7.1% of GR24 remained after incubation with SGP-7^S116A^, a value not significantly different from the substrate-only control (**Fig. 5E**). Hydrolysis of GR24, like that of YLG, therefore requires Ser116. The assay measures substrate disappearance only; the hydrolysis products were not detected or identified, and no claim is made regarding product identity.

Incubation of SGP-7 with GR24 also produced a time-dependent increase in a modified form of the protein carrying an additional 114 Da (**Supplementary Figure S8B**), rising from approximately 2.5% of the total at 15 min to approximately 17.5% at 360 min (**Fig. 5F**), accompanied by a corresponding reduction in the unmodified 34.5-kDa species to approximately 85% and 59% of the untreated control at the same two time points (**Fig. 5G**). The chemical nature of the 114-Da modification, and the identity of the modified residue, remain to be determined.

As additional structural support for the observed hydrolytic activity, the stability of the SGP-7– GR24 complex was evaluated during a 100 ns molecular dynamics simulation. The GR24 ligand RMSD remained within a narrow range throughout the simulation, with transient fluctuations but no progressive increase, indicating that the ligand remained stably accommodated within the binding pocket (**Supplementary Figure S5C**). The minimum distance between the Ser116 Oγ atom and GR24 averaged 0.31 nm and remained below 0.4 nm in 99.99% of the 50,001 frames analysed, indicating persistent close proximity of the catalytic nucleophile to the bound ligand (**Fig. 5B**). These observations provide structural support for, but do not independently demonstrate, the experimentally established hydrolytic activity of SGP-7 toward GR24.

### 2.8. Transient expression of SGP-7 in *N. benthamiana* reduces stature and releases axillary buds

Having established that SGP-7 is an active hydrolase of strigolactone analogues *in vitro*, its biological activity was assessed in a plant host, with particular attention to whether it reproduces the architectural changes reported for its orthologue RY378 in rice (Zhang et al., 2024). SGP-7 was expressed transiently in *N. benthamiana* using the potato virus X (PVX)-based vector pGR107, with the empty PVX vector as control, and plants were scored 21 days after agroinfiltration.

Plants expressing SGP-7 were markedly shorter than empty-vector controls (6.5 ± 1.3 cm versus 12.1 ± 2.3 cm; a 46% reduction; *n* = 11 plants per treatment; *P* < 0.0001) (**Fig. 4B–D**) and developed significantly more axillary buds exceeding 2 cm in length (3.5 ± 2.6 versus 0.9 ± 1.2 buds per plant; *n* = 11; *P* < 0.01) (**Fig. 4B, 4E**), indicating a release of apical dominance. Consistent with this phenotype, transcript levels of the strigolactone-responsive marker gene *NbBRC1*, measured in axillary bud tissue harvested at 21 days post-infiltration, were approximately fourfold lower than in controls (ΔΔ*C*_t_ = 2.11; fold change 0.23; *n* = 3 biological replicates; *P* < 0.01) (**Fig. 4F**).

Successful and specific expression of SGP-7 was confirmed by RT-qPCR: SGP-7 transcripts were detected only in plants infiltrated with the SGP-7 construct and were undetectable in empty-vector controls, whereas PVX coat protein transcripts were detected in both treatments (**Supplementary Figure S6B**), confirming systemic infection in both groups and allowing the comparison to isolate the effect of the effector from that of viral infection. Transient expression of an SGP-7–mGFP fusion produced fluorescence in both the nucleus and the cytoplasm, a distribution indistinguishable from that of free mGFP (**Fig. 4A**; **Supplementary Figure S6A**); given the size of the fusion protein, this nucleocytoplasmic distribution is reported as an observation and is not interpreted as evidence of targeted nuclear localisation.

These observations reproduce, in a dicot host and following transient expression, the reduced stature and increased shoot proliferation reported for RY378 in stably transformed rice and in *N. benthamiana* (Zhang et al., 2024), and are consistent with the two orthologues acting on the same developmental pathway.

## 3. Discussion

The developmental symptoms of phytoplasma disease –witches’ broom, excessive tillering, loss of apical dominance –have long suggested interference with the hormonal control of shoot architecture, and strigolactone signaling has been the obvious candidate. Until now, however, every phytoplasma effector characterized at the molecular level has been shown to act by binding and destabilizing a host protein. The description of RY378 sharpened the question considerably: that effector produces severe tillering in rice and alters strigolactone– and auxin-responsive transcripts, and its predicted α/β-hydrolase domain was proposed as the basis of its activity, but the proposal was never tested experimentally (Zhang et al., 2024). We show here that the orthologous protein SGP-7 is a genuine, catalytically active esterase, that it hydrolyses two structurally distinct strigolactone analogues, and that this activity depends on a single active-site serine. To our knowledge this is the first demonstration that a phytoplasma effector possesses enzymatic activity toward strigolactone-like molecules, and it identifies a mode of host manipulation that is mechanistically distinct from the protein-degradation strategies described for this genus.

### 3.1. A conserved effector family rather than a lineage-specific innovation

The comparative analysis revises an important aspect of the earlier description. Zhang and colleagues detected RY378 homologues only in SCWL1 and concluded that the effector is confined to the 16SrXI group and may represent a unique pathogenic mechanism (Zhang et al., 2024). Searching a purpose-built secretome database constructed from 288 phytoplasma genomes identified homologues of SGP-7 in at least seven 16Sr groups (16SrI, II, III, VII, X, XI and XIV). The discrepancy is most simply explained by database scope: homology searches restricted to annotated reference proteomes can miss effectors in draft genomes and proteins annotated as hypothetical, both of which are common features of phytoplasma genome resources. Rather than a strain-specific protein, this α/β-hydrolase therefore represents a conserved phytoplasma effector family, with homologues retained across lineages that infect both monocot and dicot hosts. The retention of a catalytically competent enzyme across these lineages, despite the extensive reductive evolution and loss of biosynthetic capacity characteristic of phytoplasma genomes, is consistent with functional importance.

### 3.2. SGP-7 is an esterase whose catalytic behavior resembles a receptor rather than a metabolic enzyme

The biochemical characterisation places SGP-7 within the serine hydrolase class. Activity declines monotonically across the *p*-nitrophenyl C2–C12 series, consistent with an esterase preference for short-chain substrates, while substitution of Ser116 abolishes activity and treatment with PMSF reduces it. The catalytic triad assignment is supported by sequence, by the position of each residue in the predicted fold, and by the presence of Ser116 and His261 within the principal surface cavity.

The kinetic parameters are among the most informative results of this study. With an apparent *K*_m_ of 3.55 µM for YLG and a *k*_cat_ of 9.38 × 10⁻³ s⁻¹, SGP-7 is a slow-turnover enzyme by metabolic standards, converting on the order of one substrate molecule every two minutes. The plant strigolactone receptors that SGP-7 structurally resembles are themselves extremely slow, and this is consistent with their function: D14 is a dual-function protein in which hydrolysis is part of hormone perception rather than simply a route to substrate disposal, and the covalently linked intermediate generated by D-ring cleavage drives the conformational change that permits MAX2 recruitment. Slow, near-single-turnover catalysis is therefore characteristic of a hydrolase functioning as a receptor. The *K*_m_ of SGP-7 is within twofold of that reported for AtD14, while its turnover number is similarly low, suggesting that SGP-7 retains aspects of the substrate-recognition properties of the receptor family rather than exhibiting the kinetics expected of an efficient catabolic enzyme.

This interpretation is reinforced by the structural comparison. CASTp analysis gives the principal SGP-7 cavity a solvent-accessible volume of 167.3 Å³, essentially identical to the 166.9 Å³ obtained for AtD14 under the same analysis, although the SGP-7 pocket is more enclosed. A bacterial protein with a cavity of strigolactone-appropriate dimensions, an equivalent catalytic triad and the ability to hydrolyze strigolactone analogues with receptor-like kinetics is consistent with a functional mimic of the host receptor.

### 3.3. A mechanistic hypothesis: hydrolysis without signal transduction

What would such a mimic do to a plant? In the host, perception and inactivation of strigolactone are inseparable: D14 hydrolyses the hormone, but the hydrolysis event is productive because it licenses assembly of the SCF^MAX2^ complex and consequent degradation of the SMXL6/7/8 repressors, which in turn de-represses *BRC1* and reinforces bud dormancy (Guercio et al., 2023). Signal output therefore requires both the chemistry and the protein–protein interface.

SGP-7 possesses the catalytic activity, but there is no evidence that it possesses the corresponding signaling interface. It is a bacterial protein that falls within a phytoplasma-specific clade clearly separated from the characterized α/β-hydrolase families, and nothing in its sequence or predicted structure indicates a surface capable of engaging MAX2. On this basis, a hypothesis consistent with, but not proven by, the present data is that SGP-7 functions as a catalytically competent but signaling-incompetent mimic of the strigolactone receptor: it consumes strigolactone-like molecules without transducing the signal that their consumption would normally generate. Such an activity could attenuate strigolactone signaling in the vicinity of the pathogen and, through reduced SMXL turnover, would be expected to leave *BRC1* transcript levels low and axillary buds released from dormancy. This is a testable proposition: a direct interaction test between SGP-7 and MAX2 or its *N. benthamiana* orthologue, with AtD14 as a positive control, would discriminate between the alternatives. That experiment was not performed here, and the model should therefore be regarded as a framework for interpretation rather than a demonstrated mechanism.

A second, non-exclusive possibility also deserves consideration. SGP-7 is most active toward the shortest acyl esters tested, and the methyl esters of several plant hormones, including methyl salicylate, methyl jasmonate and methyl indole-3-acetate, are hydrolyzed by plant methyl-esterases that belong to the same α/β-hydrolase superfamily and carry Ser–Asp–His triads with comparable spacing. These substrates were not tested and no claim is made regarding their hydrolysis; the observed substrate preference simply means that this possibility cannot presently be excluded, and it is noted as a direction for future work.

### 3.4. Partial inhibition by PMSF and the genetic evidence for Ser116

PMSF reduced SGP-7 activity by 31–33%, whereas substitution of Ser116 abolished activity entirely. The two results are not in conflict, but the partial inhibition should be interpreted with care. PMSF is unstable in aqueous buffer at neutral to slightly alkaline pH, and inhibitor decay over the course of an assay will always reduce apparent inhibition. In this experiment, however, the inhibitor was present at 1 mM against an enzyme concentration of 67 nM, and the substrate was also present at 1 mM, so PMSF and substrate were equimolar and the substrate itself acts as a competing electrophile for the active-site serine. Endpoint inhibition measured under these conditions therefore reflects the combined influence of inhibitor availability and substrate competition, and should not be read as a quantitative measure of the fraction of activity that is serine-dependent.

The genetic evidence is unambiguous. SGP-7^S116A^ expresses, purifies and elutes from size-exclusion chromatography indistinguishably from the wild-type protein, indicating that the substitution does not perturb the fold, yet it is devoid of activity against every substrate tested. We therefore regard the mutant, rather than the inhibitor experiment, as the definitive demonstration that Ser116 is the catalytic nucleophile, and report the PMSF result as corroborative chemical evidence for a serine hydrolase mechanism. A concentration series using freshly prepared inhibitor would resolve the quantitative behaviour of this reagent against SGP-7 and is a straightforward addition should it be requested.

### 3.5. GR24-dependent covalent modification of SGP-7

Treatment of SGP-7 with GR24 results in the formation of a species carrying an additional 114 Da, indicating that the protein undergoes a GR24-dependent covalent modification. The functional significance and precise chemistry of this modification remain to be established. The abundance of the modified species increased from approximately 2.5% after 15 min to approximately 17.5% after 6 h, indicating that adduct formation is a relatively slow and limited process under the conditions examined. Such a modification could be associated with the biochemical function of SGP-7, for example as part of a substrate-processing or catalytic mechanism; however, the present data do not establish whether the modified species represents a catalytic intermediate, a stable reaction product or a secondary modification. Identification of the modified residue and further characterisation of the reaction will be necessary to determine the molecular basis and functional relevance of the adduct, and to establish whether it is directly linked to SGP-7 activity.

### 3.6. Molecular dynamics supports stable association of GR24 with SGP-7

Molecular docking of GR24 into SGP-7 yielded a docking score of −5.7, with GR24 positioned within the predicted binding pocket. Residues located within 3.0 Å of the docked ligand included Ser116 and His261, both of which were also identified within the pocket predicted by CASTp, supporting the placement of GR24 within the predicted binding site. During the 100 ns simulation, the backbone RMSD of both apo SGP-7 and the GR24-bound protein remained broadly stable, with the GR24-bound form showing somewhat larger fluctuations, indicating ligand-associated changes in protein motion without evidence of overall structural destabilisation. The ligand RMSD likewise remained within a narrow range with no progressive increase, indicating that GR24 remained associated with SGP-7 throughout. No substantial differences were observed in radius of gyration or solvent-accessible surface area between the apo and GR24-bound forms, indicating that ligand binding did not appreciably change the overall size or solvent exposure of the protein. A single 100 ns trajectory cannot address longer-timescale conformational change, and the simulation is presented as a stability check rather than as evidence from independent replicate simulations; within that limit, the consistent RMSD, radius of gyration and solvent-accessible surface area profiles indicate that the docked complex remained structurally stable. The predicted involvement of Ser116 is supported experimentally by the loss of activity in SGP-7^S116A^. Taken together, the docking, cavity analysis, molecular dynamics and mutational data provide complementary evidence for the predicted GR24-binding site and for the functional contribution of Ser116.

### 3.7. A distinct strategy within the phytoplasma effector repertoire

The mode of action described here differs fundamentally from that of the other characterized phytoplasma effectors. SAP11 destabilizes TCP transcription factors (Pecher et al., 2019), SAP54/PHYL1 directs MADS-box proteins for degradation (MacLean et al., 2011), SAP05 subverts the proteasomal Rpn10 subunit to eliminate SPL and GATA factors (Huang et al., 2021), and SWP1 destabilises BRC1 protein (N. Wang et al., 2018). Each acts stoichiometrically on a host protein through a binding interface. SGP-7 instead acts catalytically on a small molecule. This distinction has a practical corollary for how such effectors are detected: an effector that functions enzymatically need not accumulate to high levels or engage an abundant host target, and could therefore be missed by the interaction-based screens through which most phytoplasma effectors have been discovered.

The contrast with SWP1 is particularly instructive, because both effectors converge on the same developmental output through different points in the same pathway. SWP1 acts downstream, removing BRC1 protein directly (N. Wang et al., 2018). SGP-7, on the present evidence, acts upstream, on the hormonal input that determines *BRC1* transcription. That two phytoplasma lineages should have arrived at witches’ broom by such different routes underlines the importance of bud-dormancy release to phytoplasma biology –an unsurprising feature for a phloem-limited pathogen that depends on new vegetative growth and on the insect vectors that such growth attracts.

### 3.8. Interpretation of the *in-planta* results

Transient expression of SGP-7 in *N. benthamiana* reduced stature, released axillary buds and lowered *NbBRC1* transcript levels. These observations reproduce, in a dicot host and with an orthologous protein, the architectural phenotype reported for RY378 in stably transformed rice, supporting a conserved effect of this effector family on shoot architecture across host classes. The reduction in *NbBRC1*, a strigolactone-responsive marker, is consistent with attenuation of strigolactone signaling. *BRC1* integrates multiple hormonal and environmental signals, and strigolactone can also regulate branching through BRC1-independent pathways, so the present results establish association rather than causation; within that limit they provide *in planta* support for a link between SGP-7 activity and strigolactone-regulated shoot architecture. The PVX-based transient system in a heterologous dicot host provides a relevant functional model, and the relationship between catalytic activity and phenotype can be resolved directly by expressing the S116A mutant in planta. Finally, the biochemical data establish activity against synthetic strigolactone analogues; activity toward native strigolactones remains to be tested and is a defined objective for future work.

## 4. Conclusions

SGP-7, a broadly conserved phytoplasma effector, is a catalytically active esterase that hydrolyses the strigolactone analogues YLG and *rac*-GR24 in a manner strictly dependent on Ser116, with kinetic parameters and a substrate cavity comparable to those of plant strigolactone receptors. These findings supply the biochemical function that was hypothesized but left untested for this effector class, and show that the family is distributed across at least seven phytoplasma groups rather than confined to a single lineage. Transient expression in *N. benthamiana* reduces stature, releases axillary buds and lowers a strigolactone-responsive marker transcript, reproducing in a dicot the architectural phenotype reported for RY378 in rice. Together, these findings identify enzymatic modification of a host hormone as a mode of action within the phytoplasma effector repertoire, and provide a specific, testable model of catalytically competent but signaling-incompetent receptor mimicry for how this widespread effector family may contribute to the developmental symptoms that characterize phytoplasma disease.

## 5. Materials and Methods

### 5.1. Genome sequencing, assembly and annotation

DNA was extracted from symptomatic sugarcane stem tissue using a modified CTAB method (Doyle et al., 1987), and phytoplasma identity and copy number were established by Sanger sequencing and quantitative PCR respectively. Long-read sequencing was performed on an Oxford Nanopore FLO-MIN114 flow cell, generating 9.71 Gb across 5.75 million reads. Raw POD5 files were base-called with Dorado (Oxford Nanopore Technologies, 2023) and reads were quality-filtered at a minimum Q-score of 20 using NanoFilt v2.8.0 (De Coster et al., 2018). *De novo* assembly with Flye v2.9.4 (Kolmogorov et al., 2020) produced 16,990 contigs, which were binned with MetaBAT2 (Kang et al., 2019); taxonomic classification with CheckM v1.2.4 (Parks et al., 2015) identified a phytoplasma bin comprising two contigs. Previously generated Illumina HiSeq data for the same sample (22,765,620 paired-end reads) were quality-assessed with FastQC v0.11.8 (Andrews, 2010), mapped to the phytoplasma bin and used for polishing with Pilon (Walker et al., 2014) (**Supplementary Figure S1**). Genome completeness was assessed with BUSCO using the phytoplasma database (Manni et al., 2021) (198 of 202 complete BUSCOs; 98%) and independently with CheckM v1.2.4 using the ‘*Candidatus* Phytoplasma’ marker set on the PGAP gene set (94.18% completeness, 0.58% contamination). The assembly was annotated using the NCBI Prokaryotic Genome Annotation Pipeline (Tatusova et al., 2016) and Prokka (Seemann, 2014), and coding sequences were assigned to KEGG (Kanehisa et al., 2016) and COG (Tatusov, 2000) categories for metabolic reconstruction.

### 5.2. Pan-genome analysis

Pan-genome analysis was performed in anvi’o v9 (Eren et al., 2020) using the updated ‘*Ca.* P. sacchari’ SCGS assembly together with 11 complete phytoplasma reference genomes retrieved from the NCBI RefSeq database. A contigs database was generated for each assembly, conserved single-copy genes were identified, and predicted protein-coding genes were functionally annotated against COG and KEGG within the anvi’o environment. Gene clusters were defined using the Markov cluster algorithm and classified as core, accessory or strain-specific according to their distribution across genomes.

### 5.3. Effector prediction and homology searching

The 461 predicted SCGS proteins were screened for N-terminal signal peptides using SignalP 6.0 (Teufel et al., 2022) and for transmembrane helices using TMHMM 2.0 (Krogh et al., 2001). Proteins with transmembrane domains outside the predicted signal peptide region were excluded, and the remainder retained as candidate secreted effectors. Subcellular targeting was predicted with LOCALIZER v1.0.4 (Sperschneider et al., 2017).

To assess distribution across phytoplasma lineages, a reference secretome database was constructed from 288 phytoplasma genomes available in NCBI. Proteins were screened with SignalP v6.0, yielding 4,838 non-redundant SignalP-positive sequences, and transmembrane helices in the mature region were predicted with TMHMM v2.0. Proteins were classified into four categories according to the relationship between the SignalP cleavage site and TMHMM predictions: Class A, no predicted transmembrane helix (*n* = 771); Class B, a single helix wholly within the signal peptide region (*n* = 3,105); Class C, a single helix extending beyond the predicted cleavage site (*n* = 662); and Class D, more than one predicted helix, considered membrane-associated (*n* = 300). Of the Class C proteins, 631 showed transmembrane extensions of ≤8 amino acids beyond the cleavage site, a discrepancy attributable to uncertainty in boundary prediction rather than to membrane localization; these were retained. The resulting relaxed secretome comprised Classes A and B together with the 631 borderline Class C proteins, giving 4,507 candidate secreted proteins. SCGS effector candidates were searched against this database using BLASTP with thresholds of ≥30% sequence identity, ≥50% query coverage and *E* ≤ 1 × 10⁻⁵, and homologue distribution was summarized by 16Sr group.

### 5.4. Structure prediction, cavity analysis, docking and molecular dynamics

The mature SGP-7 sequence (signal peptide removed) was submitted to the AlphaFold Server (Abramson et al., 2024) and the resulting models assessed with RAMPLOT (Kumar & Rathore, 2025); model 1 was selected for all downstream analyses. Surface cavities were identified with CASTp 3.0 (Tian et al., 2018); the equivalent analysis was applied under identical probe radius and settings to the *Arabidopsis* strigolactone receptor AtD14 (PDB 4IH4), and solvent-accessible (SA) area and volume are reported throughout.

Model 1 of SGP-7 was used for molecular docking of GR24 using the Schrödinger molecular modelling suite. Molecular docking of GR24 with the predicted SGP-7 structure was performed using the Glide module of Schrödinger Suite (Glide 2025-3, build 131). Docking was performed using the Standard Precision (SP) protocol with the OPLS_2005 force field, and poses were ranked according to the GlideScore. The highest ranked SGP-7–GR24 docked complex was used for molecular dynamics simulation, with apo SGP-7 simulated in parallel as a reference system under identical conditions. All-atom simulations were performed using GROMACS v2021.3 (Abraham et al., 2015) with the OPLS-AA force field. The protein was centered in a cubic box with a minimum protein–box distance of 1.0 nm and solvated using the SPC/E water model. GR24 was represented using OPLS-AA parameters generated by LigParGen. The systems were neutralized with Na⁺ and Cl⁻ ions introduced by replacing solvent molecules. Energy minimization was performed using the steepest-descent algorithm, followed by 100 ps NVT equilibration at 300 K and 100 ps NPT equilibration at 300 K and 1.0 bar, with position restraints maintained on protein and ligand during equilibration. Production simulations were performed for 100 ns (50,000,000 steps; 2 fs time step) in the NPT ensemble at 300 K and 1.0 bar with position restraints removed. One simulation was performed per system. Trajectories were corrected for periodic boundary effects and fitted to remove rotational and translational motion prior to analysis. Protein backbone RMSD, per-residue RMSF, radius of gyration, solvent-accessible surface area, ligand RMSD and the minimum distance between the Ser116 Oγ atom and GR24 were calculated from the resulting trajectories.

### 5.5. Phylogenetic analysis

The top 30 BLAST hits from the ESTHER database (Lenfant et al., 2012) were combined with the top 20 hits from a manually curated phytoplasma database comprising proteins from 19 RefSeq reference genomes. Sequences were aligned with MAFFT v7.2.5 (Katoh, 2002) and a maximum-likelihood tree inferred with IQ-TREE v3.1.2 (Wong et al., 2026) under the Q.PFAM+R5 substitution model with 1,000 bootstrap replicates.

### 5.6. Alkaline phosphatase (PhoA) secretion assay

Assays were performed essentially as described (Liu et al., 2019). The full-length *E. coli* K12 *phoA* gene and a derivative lacking its native signal peptide (*mphoA*) were cloned into pET21a using NdeI and XhoI as positive and negative controls respectively. The predicted SGP-7 signal peptide was synthesized and fused in frame to *mphoA* in pET21a, PCR amplification of this region having been unsuccessful. Constructs were transformed into chemically competent *E. coli* BL21(DE3) and transformants selected on LB agar supplemented with 5-bromo-4-chloro-3-indolyl phosphate (90 µg mL⁻¹), ampicillin (50 µg mL⁻¹), 0.1 mM IPTG and Na₂HPO₄ (75 mM), and incubated at 37 °C for 18 h. Because PhoA is enzymatically active only after translocation to the periplasm, export mediated by a functional signal peptide produces blue colonies on BCIP, whereas white colonies indicate the absence of export.

### 5.7. Protein expression and purification

The SGP-7 sequence lacking the signal peptide was codon-optimized for *E. coli* and cloned into pET21a under the T7 promoter with a C-terminal hexa-histidine tag using NdeI and XhoI. The S116A substitution was introduced using the Phusion Site-Directed Mutagenesis Kit (Thermo Fisher, F541) and verified by Sanger sequencing. Constructs were transformed into *E. coli* BL21(DE3).

Overnight primary cultures (10 mL LB, 100 µg mL⁻¹ ampicillin, 37 °C, 220 rpm) were used to inoculate 400 mL LB containing 100 µg mL⁻¹ ampicillin, grown at 37 °C and 180 rpm for 2 h. Expression was induced with 0.28 mM IPTG and cultures incubated at 18 °C and 180 rpm for 24 h. Cells were harvested at 4,500 × *g* for 15 min at 4 °C, resuspended in lysis buffer (50 mM Tris-HCl pH 8.0, 300 mM NaCl, 0.1% NP-40, 5% glycerol) with protease inhibitor cocktail, incubated on ice for 30 min and disrupted by sonication (70% amplitude, 2 s on/10 s off, 1 min total). Lysates were clarified at 13,500 rpm for 45 min at 4 °C.

Ni-NTA resin (4 mL) was equilibrated with deionized water followed by equilibration buffer, and the cleared supernatant was incubated with resin at 4 °C with gentle rotation for 5 h. Resin was transferred to gravity-flow columns, washed sequentially with lysis buffer and wash buffer (50 mM Tris-HCl pH 8.0, 300 mM NaCl, 5% glycerol, 0.1% NP-40, 30 mM imidazole), and bound protein eluted in 15 mL elution buffer (50 mM Tris-HCl pH 8.0, 300 mM NaCl, 5% glycerol, 300 mM imidazole, protease inhibitor cocktail) collected as 2 mL fractions. Fractions were analysed by SDS-PAGE, pooled and concentrated using Amicon Ultra centrifugal filters at 2,500 × *g* and 4 °C. Concentrated protein was applied to a Superdex 200 Increase 10/300 GL column (Cytiva) on an ÄKTA system pre-equilibrated in 1× PBS (pH 7.4), run at 0.2 mL min⁻¹ with detection at 280 nm, simultaneously removing imidazole and exchanging the protein into PBS. Fractions corresponding to the major peak (16–18 mL) were pooled. Protein concentration was determined by BCA assay.

### 5.8. Immunoblotting and mass spectrometry

Purified proteins were separated by SDS-PAGE, transferred to PVDF, probed with a mouse anti-6×His primary antibody (1:1000) followed by an HRP-conjugated anti-mouse secondary antibody, and detected by chemiluminescence. Intact mass determination was performed on a ZenoTOF 7600 mass spectrometer (SCIEX) coupled to a Nexera 40 Series UHPLC (Shimadzu), with separation on a ZORBAX 300SB-C8 column (4.6 × 150 mm, 5 µm; Agilent) in positive-ion mode; the observed mass was compared with that predicted from the amino acid sequence of the tagged construct. For peptide-level identification, the protein was digested with trypsin and the resulting peptides analysed by LC-MS/MS, with data searched using MaxQuant. Intact and peptide mass spectrometry were performed on wild-type SGP-7 only.

### 5.9. Esterase activity assays with *p*-nitrophenyl esters

Stock solutions (50 mM) of *p*-nitrophenyl acetate (C2), butyrate (C4), octanoate (C8), decanoate (C10) and dodecanoate (C12) were prepared in acetonitrile, stored at −20 °C, and diluted to 1 mM in PBS (pH 7.4) before use. Reactions (100 µL) contained 67 nM purified SGP-7, 100 µM substrate and PBS, with the final acetonitrile concentration maintained at 1% (v/v). Plates were incubated at 30 °C for 1 h and *p*-nitrophenol release monitored at 405 nm on a Multiskan SkyHigh spectrophotometer (Thermo Fisher Scientific, USA). Substrate-only controls without enzyme were included to correct for non-enzymatic hydrolysis and background absorbance. Background-corrected absorbance was converted to *p*-nitrophenol concentration using a standard curve prepared in PBS containing 1% acetonitrile, and activity expressed as specific activity (U mg⁻¹, where one-unit releases 1 µmol *p*-nitrophenol per minute). Relative activity was calculated within each assay by setting the highest specific activity (C2) to 100%. Each assay used four technical replicates and was performed in three independent assays.

### 5.10. Time-course hydrolysis

Reactions (100 µL) contained 100 µM C2 or C4 substrate in 1× PBS (pH 7.4) with 1% (v/v) acetonitrile and 0.13 µM SGP-7 or SGP-7^S116A^. Absorbance at 405 nm was recorded at 1-min intervals for 1 h at 30 °C with continuous shaking between reads, and converted to *p*-nitrophenol concentration using the standard curve. Each point represents the mean of eight technical replicates.

### 5.11. PMSF inhibition

Reactions (100 µL) contained 1× PBS (pH 7.4), 1 mM substrate (C2 or C4), 67 nM SGP-7 and 1 mM phenylmethylsulfonyl fluoride, and were incubated for 30 min at 30 °C. Control reactions lacking PMSF were performed under identical conditions. *p*-Nitrophenol release was measured at 405 nm and quantified against the standard curve. Experiments were performed in three independent assays with four technical replicates each.

### 5.12. YLG hydrolysis and kinetic analysis

Assays were performed in black 96-well microplates in 100 µL containing purified SGP-7 or SGP-7^S116A^ at 0.067 µM, 1× PBS (pH 7.4) and 1 µM Yoshimulactone Green, with DMSO maintained at 1%. Plates were shaken for 30 s before the initial and each subsequent read, and fluorescence monitored on a Mithras LB 943 multimode plate reader (excitation 480 nm, emission 520 nm) at 60 s intervals for approximately 2 h at 25 °C. Background fluorescence from no-enzyme controls was subtracted. The time course used eight technical replicates.

For kinetic analysis, assays were performed as above at YLG concentrations of 1, 2.5, 5, 10, 15, 20 and 40 µM, with fluorescence recorded at 60 s intervals for approximately 1 h. Background fluorescence from no-enzyme controls at each substrate concentration was subtracted and fluorescence converted to fluorescein concentration using a fluorescein standard curve. Initial velocities were determined by linear regression over the initial linear phase (first 402 s; *R*² > 0.98) and fitted to the Michaelis–Menten equation by nonlinear regression in GraphPad Prism v8.0.2. Experiments comprised three independent biological replicates with four technical replicates each.

### 5.13. GR24 hydrolysis and HPLC quantification

Reactions (100 µL) contained purified SGP-7 and *rac*-GR24 in PBS (pH 7.4). Racemic GR24 (a mixture of GR24^5DS^ and GR24^ent-5DS^) was used throughout. Two controls were included: GR24 alone, and GR24 with SGP-7^S116A^ under identical conditions. Reactions were incubated at 30 °C for 30 min, then extracted with 500 µL HPLC-grade ethyl acetate, vortexed and centrifuged at 14,000 rpm for 10 min. The organic phase was collected, dried under vacuum and reconstituted in 500 µL HPLC-grade methanol. Samples (50 µL) were analysed on a Shimadzu HPLC system with a reverse-phase C18 column (4.1 × 150 mm, 5 µm) using a methanol–water gradient at 0.8 mL min⁻¹ over 35 min, with methanol increased to 90% over 19 min, held for 6 min, then returned to 10% over 10 min, and photodiode-array detection at 254 nm. GR24 standards (10, 32, 50, 100, 500 and 1000 µM) were analysed under identical conditions to establish retention time, and peak areas were used to quantify remaining GR24. Values are expressed as a percentage of the substrate-only control. The experiment comprised three biological replicates with three technical replicates each. Only substrate disappearance was measured; hydrolysis products were not quantified, as authentic standards were not available.

### 5.14. Intact mass spectrometry of the SGP-7–GR24 reaction

Purified SGP-7 (32 µM) was incubated with GR24 (500 µM) in 1× PBS (100 µL) at 30 °C. Matched controls contained SGP-7 (32 µM) with 2.5% (v/v) DMSO. Samples were denatured with 10% TFE for 2 h at room temperature, centrifuged for 10 min, and 2 µL of each sample was subjected to LC-MS analysis. Intact mass analysis was performed using a ZenoTOF 7600 coupled to a Nexera 40 Series UHPLC/HPLC system with a Zorbax 300SB-C8 column (4.6 × 150 mm, 5 µm). MS data were acquired in positive-ion mode over 400–2000 Da, with an ion spray voltage of 5500 V and an accumulation time of 0.5 s. Spectra were deconvoluted, and peaks corresponding to 34,541 Da and 34,656 Da (+114 Da) were quantified. Three technical replicates were analysed at each time point.

### 5.15. Subcellular localization

*N. benthamiana* seeds were sown in a mixture of Soilrite, perlite, vermiculite and cocopeat (2:1:1:2), stratified at 4 °C in darkness for 3 d, and grown at 23 °C under a 16 h light/8 h dark photoperiod at 60% relative humidity.

The SGP-7 coding sequence, codon-optimised for *N. benthamiana*, was cloned into pEG103 in frame with a C-terminal mGFP tag; the empty pEG103-mGFP vector served as control. Constructs were transformed into *Agrobacterium tumefaciens* GV3101. Cultures were grown in LB with kanamycin (50 µg mL⁻¹), gentamicin (30 µg mL⁻¹) and rifampicin (10 µg mL⁻¹) to OD₆₀₀ 0.6, harvested, washed twice in 10 mM MgCl₂ and resuspended in 10 mM MgCl₂ to OD₆₀₀ 0.6. Acetosyringone was added to 100 µM from a freshly prepared 100 mM DMSO stock and suspensions were incubated for 1 h in darkness with gentle shaking before infiltration into leaves of 4-week-old plants using a needleless syringe. Imaging was performed 48–72 h post-infiltration on an Olympus FV3000 confocal microscope, detecting GFP fluorescence and chlorophyll autofluorescence. Expression was confirmed by immunoblotting with an anti-GFP antibody. Experiments were performed in three independent biological replicates.

### 5.16. Transient overexpression and phenotyping

The codon-optimized SGP-7 sequence was cloned into the PVX-based vector pGR107 using the SmaI site and transformed into *A. tumefaciens* GV3101 carrying the pSOUP helper plasmid. Cultures carrying either the SGP-7 construct or the empty vector were grown to OD₆₀₀ 0.6, harvested at 5,000 × *g* for 10 min and resuspended in infiltration buffer (10 mM MgCl₂, 100 µM acetosyringone). Two leaves per plant were infiltrated at the four-week stage, with 11 plants per treatment, each plant constituting one biological replicate. Plants were monitored for 21 d; plant height was measured and axillary buds longer than 2 cm were counted at 21 days post-infiltration. Statistical significance between control and test treatments was assessed using an unpaired two-tailed Student’s *t*-test for plant height and a two-tailed Mann–Whitney U test for axillary bud number.

### 5.17. RNA extraction and RT-qPCR

Axillary buds were collected from three biological replicates each of empty-vector and SGP-7-expressing plants at 21 days post-infiltration. Total RNA was extracted with the RNeasy Plant Mini Kit (Qiagen, 74904) and assessed on a NanoDrop spectrophotometer. First-strand cDNA was synthesized from 1 µg total RNA in 20 µL using the QuantiTect Reverse Transcription Kit (Qiagen, 205311), including the integrated gDNA Wipeout step, and diluted 1:5 before use. Reactions (10 µL) contained 5 µL TB Green Premix Ex Taq II (Tli RNaseH Plus; Takara Bio, RR820A), 0.1 µM of each primer and 1 µL diluted cDNA, and were run on a CFX96 Real-Time PCR Detection System (Bio-Rad) using two-step cycling (95 °C for 5 s; 60 °C for 30 s; 40 cycles), followed by melt-curve analysis to confirm amplification specificity. Each biological replicate was analysed in three technical replicates. Relative transcript abundance was calculated by the 2^−ΔΔCt^ method with *PP2A* as reference and empty-vector samples as calibrator. Statistical significance was assessed by unpaired two-tailed Student’s *t*-test applied to Δ*C*_t_ values in GraphPad Prism v8.0.2.

### 5.18. Statistical analysis

Enzymatic assays were performed in three independent assays with the technical replication stated for each assay, and are presented as mean ± SD. Unless stated otherwise, phenotypic comparisons used unpaired two-tailed Student’s *t*-tests. RT-qPCR data were analysed by unpaired two-tailed Student’s *t*-test applied to Δ*C*_t_ values, which is the appropriate approach for quantitative PCR data. All analyses were performed in GraphPad Prism v8.0.2. Significance is indicated as \**P* < 0.05, \*\**P* < 0.01, \*\*\**P* < 0.001, \*\*\*\**P* < 0.0001.

## Data availability

The improved genome assembly of ‘*Candidatus* Phytoplasma sacchari’ isolate SCGS has been deposited in the NCBI database under BioProject ID PRJNA565134 and BioSample ID SAMN12735220. The complete circular chromosome and plasmid sequences are available in GenBank under accession numbers CP176574.1 and CP176575.1 respectively.

## Acknowledgements

The authors acknowledge the University Grants Commission (UGC), Government of India, for financial support through the UGC-NET Junior Research Fellowship; Award No. 211610161705. The authors thank Dr Nripendra Verma (Institute for Stem Cell Science and Regenerative Medicine, inStem, Bengaluru) for support with the proteomics studies and for valuable suggestions, and Dr Sangeeta Sawant, Director, Bioinformatics Centre, Savitribai Phule Pune University, for valuable inputs and guidance on the molecular dynamics simulations. The authors acknowledge the Bioimaging Facility at the National Centre for Cell Science (NCCS), Pune, and thank Ashwini Atre and Trupti Kulkarni for their support. The authors acknowledge the Department of Biotechnology (DBT) for support through grant BT/PR40189/BTIS/137/50/2022, under which the Schrödinger software used for molecular docking in this study was procured.

## Author contributions

SV planned the study, performed the experiments, analysed and interpreted the data, and prepared the original draft of the manuscript. AY conceptualized and supervised the study, contributed to experimental design and data interpretation, critically revised the manuscript, and approved the final version.

