## Supplemental Data for "A broadly conserved phytoplasma α/β-hydrolase effector, SGP-7, is a catalytically active esterase that hydrolyses strigolactone analogues and alters shoot architecture"

### This PDF file includes:

- Figures S1 to S8
- Table S1

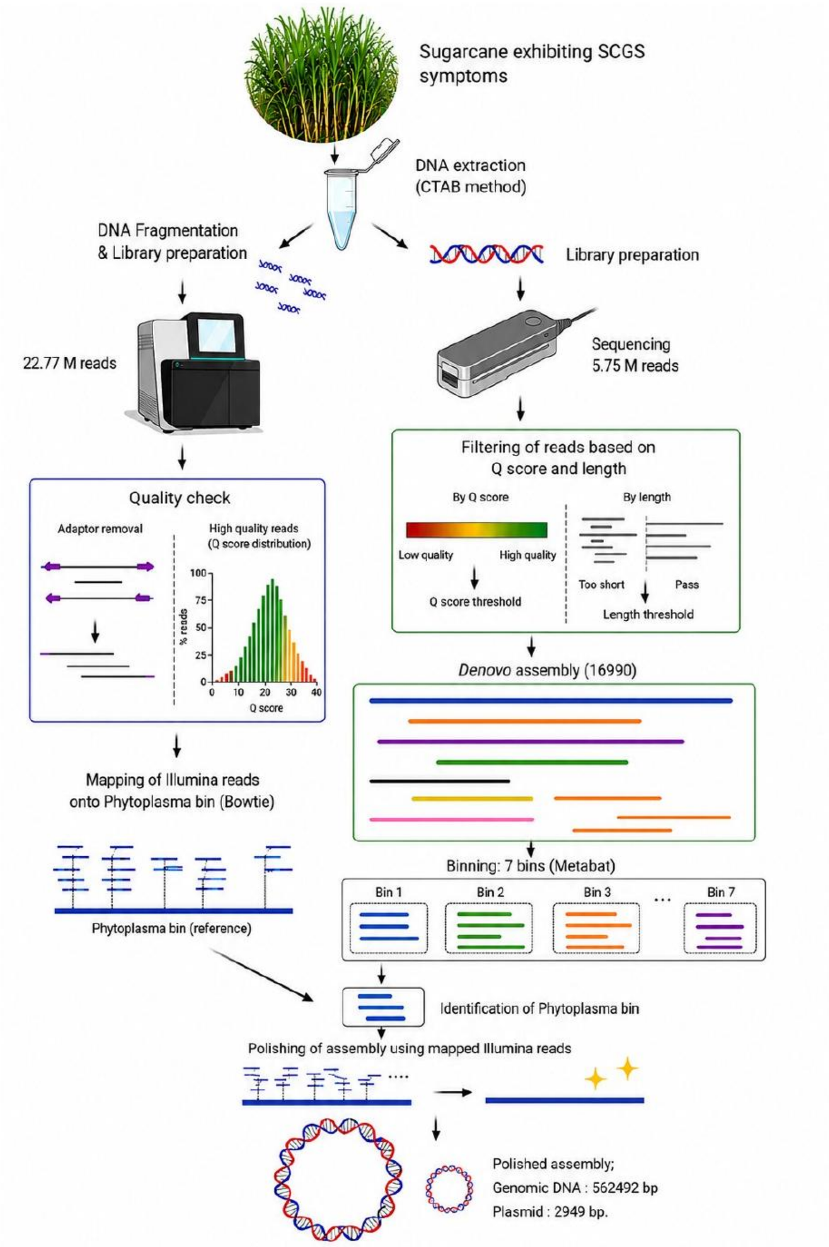


**Fig. S1.** Workflow for phytoplasma genome reconstruction from symptomatic sugarcane. Long-read sequencing, assembly, binning, taxonomic assignment and short-read polishing steps are shown in sequence.

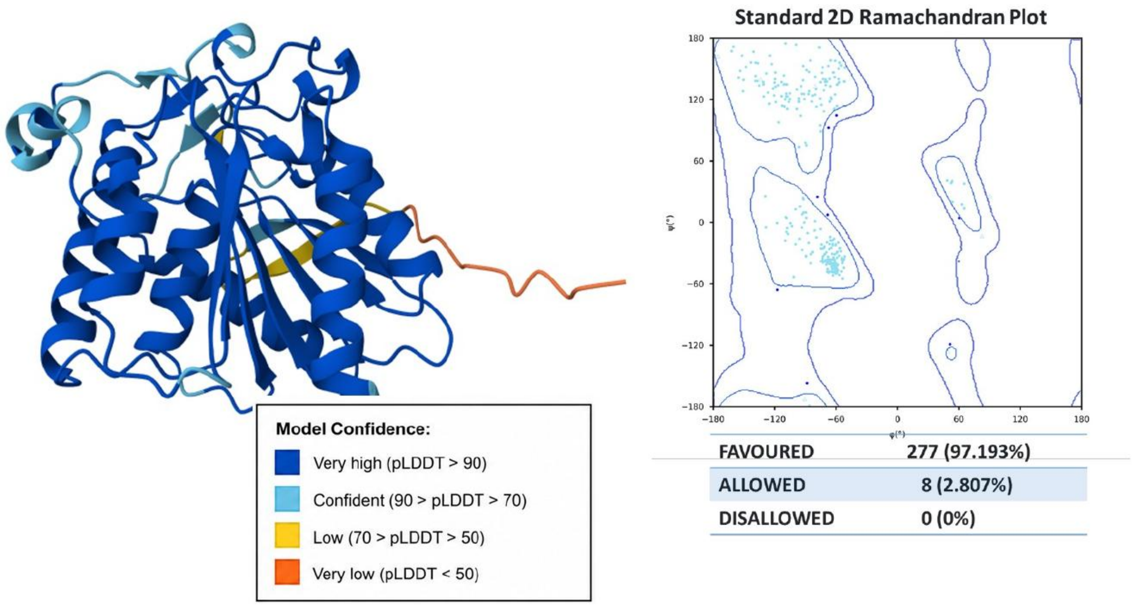


**Fig. S2.** Structural model and Ramachandran validation of SGP-7. The predicted SGP-7 structure is colored according to AlphaFold 3 per-residue confidence (pLDDT), together with the Ramachandran plot showing the distribution of residues across favored, allowed and disallowed regions.

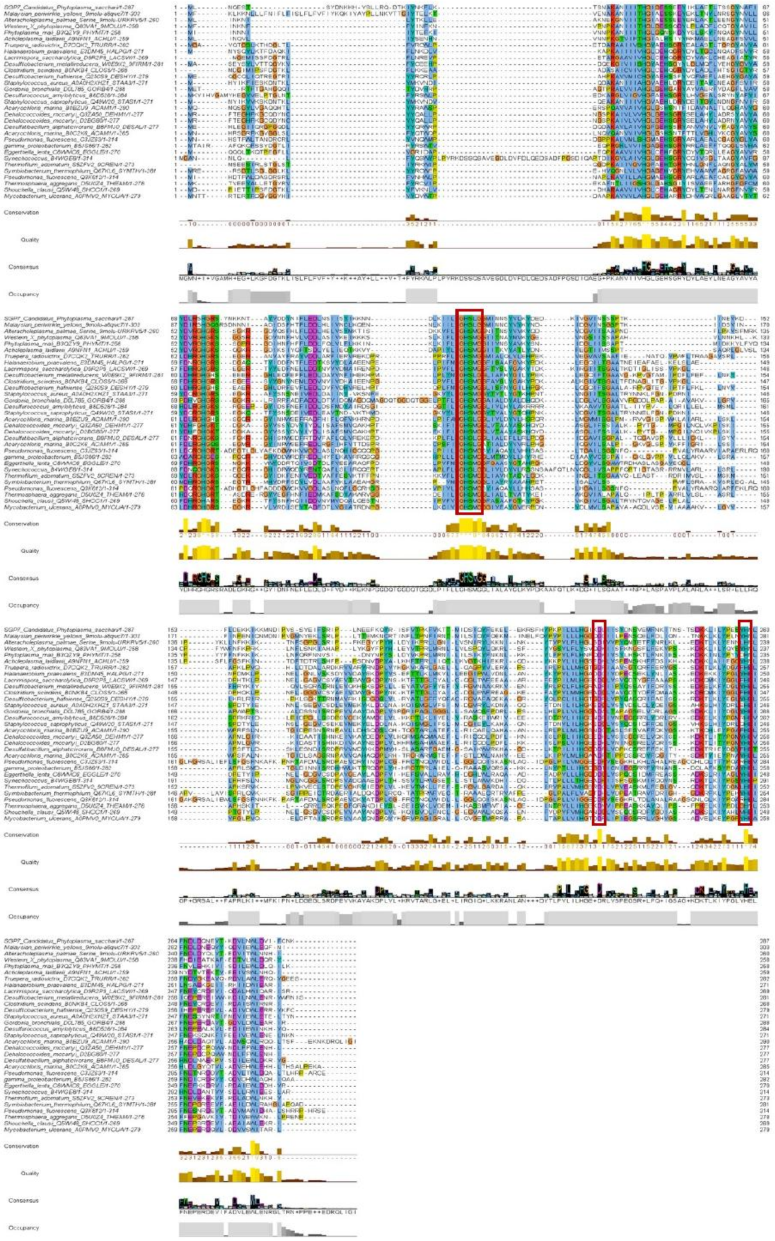

**Fig. S3.** Multiple sequence alignment of SGP-7 and homologous α/β-hydrolase proteins. Alignment was performed with MAFFT using SGP-7 as query against the top 30 homologs retrieved from the ESTHER database. Conserved residues are shaded by degree of conservation, and the catalytic triad residues Ser116, Asp229 and His261 are indicated.

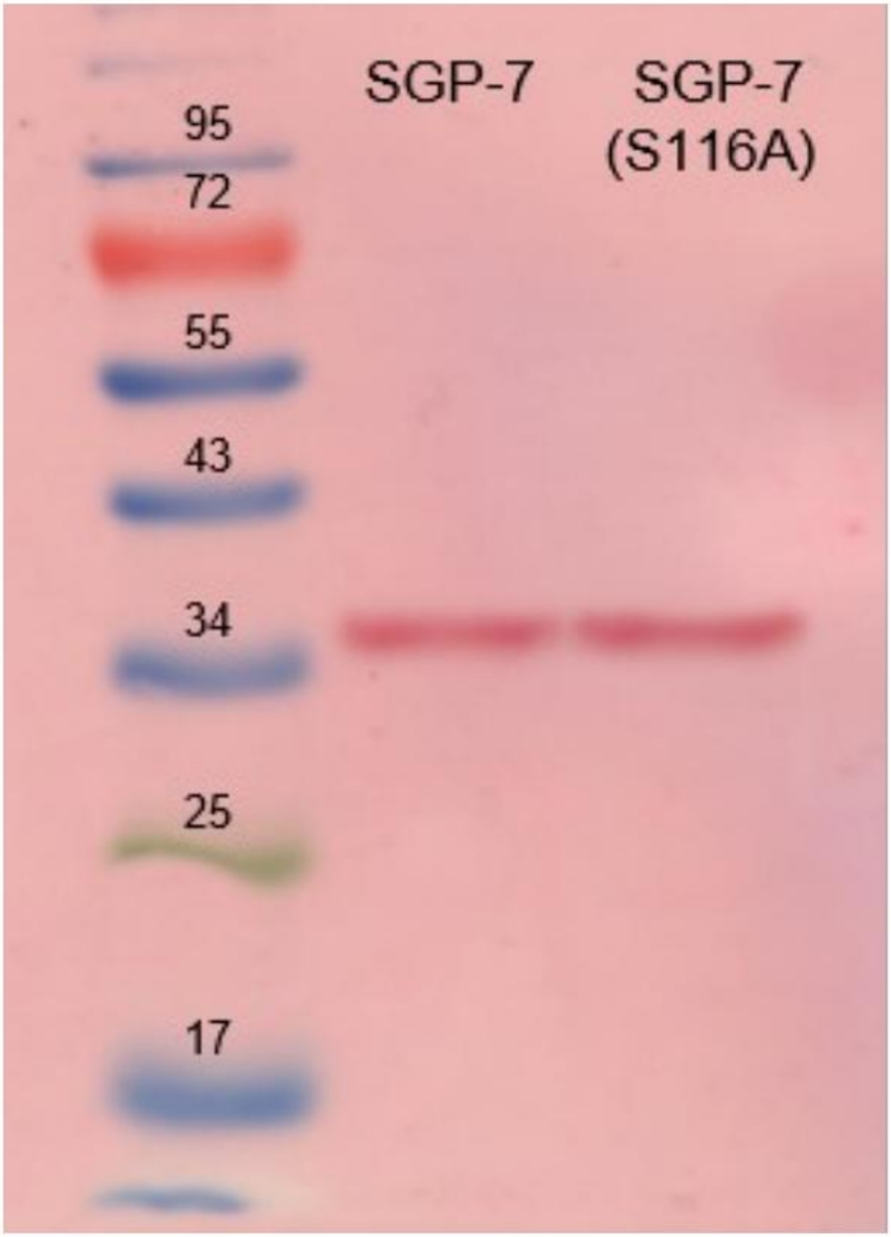


**Fig. S4.** SDS-PAGE analysis of purified SGP-7 and SGP-7$^{S116A}$. Both proteins migrate as a predominant band at approximately 34 kDa. Molecular mass markers are indicated at left.

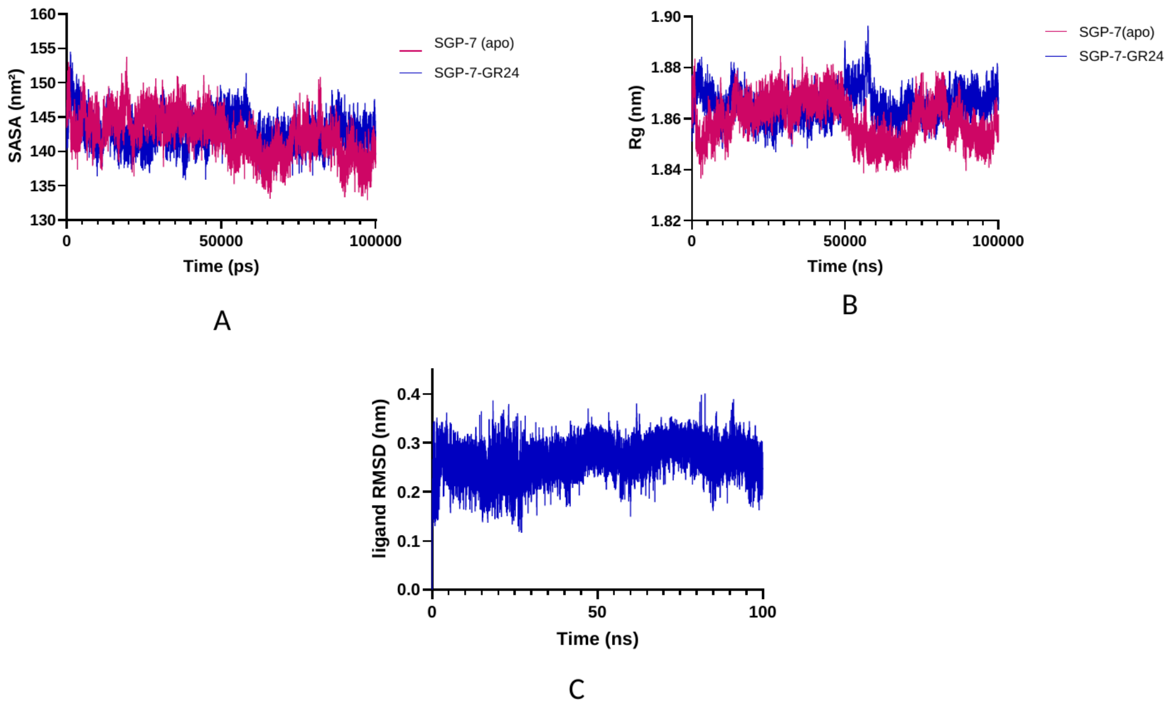


**Fig. S5.** Molecular dynamics trajectory analysis of SGP-7 and the SGP-7–GR24 complex. (*A*) Solvent-accessible surface area, (*B*) radius of gyration and (*C*) RMSD of the GR24 ligand over the 100 ns simulation. Apo SGP-7 and the SGP-7–GR24 complex are shown in magenta and blue respectively.

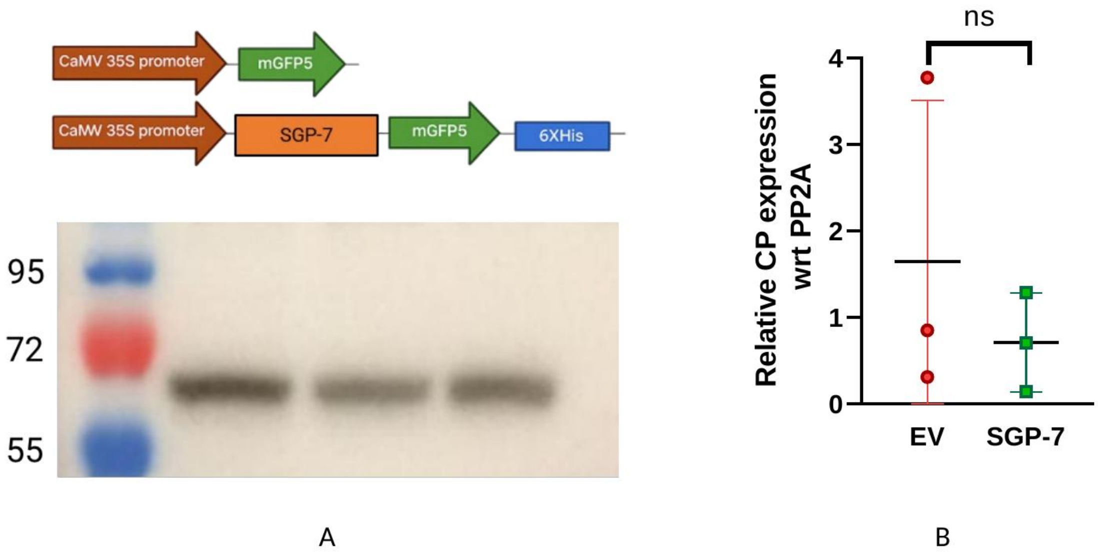


**Fig. S6.** SGP-7 expression in *Nicotiana benthamiana*. (*A*) Immunoblot confirming GFP fusion expression in plants expressing the indicated construct. (*B*) Relative expression of the potato virus X coat protein in pGR107-infiltrated plants, quantified from three biological replicates. Data are mean ± SD.

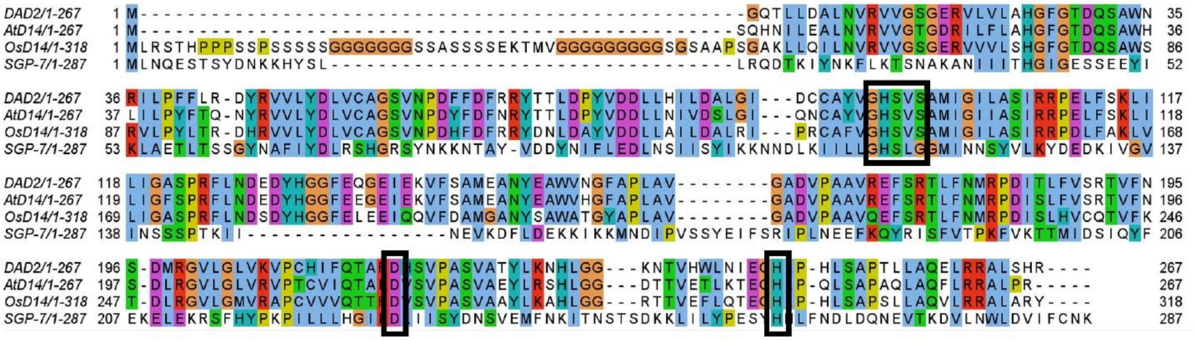


**Fig. S7.** Multiple sequence alignment of mature SGP-7 with the plant strigolactone receptors AtD14 (*Arabidopsis thaliana*), OsD14 (*Oryza sativa*) and DAD2 (*Petunia hybrida*), generated with PROMALS3D and visualized in Jalview. Boxes indicate the catalytic triad positions corresponding to Ser116, Asp229 and His261 of SGP-7.

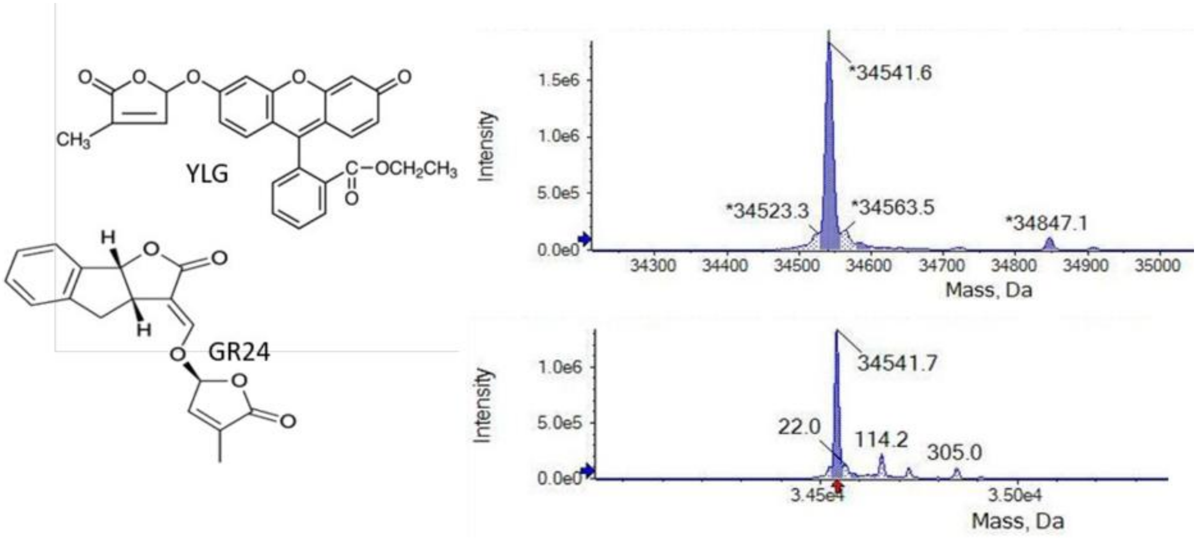


**Fig. S8.** Intact mass spectrometric analysis of SGP-7. (*Left*) Structures of the fluorogenic strigolactone analog YLG and the chemical analog GR24. (*Upper Right*) Intact mass spectrum of SGP-7 showing a major species at approximately 34,541.6 Da. (*Lower Right*) Intact mass spectrum of SGP-7 following incubation with GR24, showing an additional species at +114 Da.

**Table S1.** CASTp analysis of the predicted ligand-binding pockets of OsD14, DAD2, AtD14 and SGP-7. Solvent-accessible (SA) area ($Å^2$) and SA volume ($Å^3$) were calculated for each protein under identical probe radius and settings.

| **Protein** | **SA area ($Å^2$)** | **SA volume ($Å^3$)** |
|---|---|---|
| OsD14 | 191.575 | 100.953 |
| DAD2 | 253.499 | 90.722 |
| AtD14 | 321.575 | 166.852 |
| SGP-7 | 231.684 | 167.308 |
